# Endemic demographics of the Jomon people estimated based on complete mitogenomes reveal their regional diversity

**DOI:** 10.1101/2024.05.16.594064

**Authors:** Koki Yoshida, Yoshiki Wakiyama, Guido Valverde, Akio Tanino, Daisuke Waku, Takafumi Katsumura, Motoyuki Ogawa, Tomohito Nagaoka, Kazuaki Hirata, Kae Koganebuchi, Yusuke Watanabe, Jun Ohashi, Minoru Yoneda, Ryuzaburo Takahashi, Hiroki Oota

## Abstract

The Jomon culture that spread across Japanese archipelago began about 16,000 years ago and lasted for over 10,000 years. The genetic diversity of the Jomon people, prehistoric hunter-gatherers bearing the Jomon culture, is of great interest in understanding prehistoric East Eurasians. Until now, their demographic history has been estimated using archaeological sites and present-day genomes, but detailed studies using Jomon genomes have been insufficient. To investigate the Jomon demography, we determined the complete mitochondrial genome (mitogenome) sequences from 13 Jomon individuals and conducted population genetic analysis on 40 Jomon genomes including previously published data. Our results revealed an effective population size increase during the Incipient – Initial phase of the Jomon period, which had not been observed in analysis of mitogenome sequences from present-day Japanese populations. This endemic demographic pattern is pronounced in the eastern part of the archipelago, under the assumption of no gene flow between the Eastern and Western Jomon.

## Introduction

The Jomon people are prehistoric hunter-gatherers who inhabited the Japanese archipelago, located in the eastern part of the Eurasian continent. They are characterized by the earliest use of pottery adorned with cord-marking patterns, from which the name “Jomon” is derived, and a sedentary lifestyle, which is uncommon among hunter-gatherers (Habu, 2004). The Jomon period is divided into six phases based on temporal change in the style of pottery: Incipient (ca. 16,500-11,300 years ago (ya)), Initial (ca. 11,300-7,100 ya), Early (ca. 7,700-5,400 ya), Middle (ca. 5,400-4,400 ya), Late (ca. 4,400-3,200 ya), and Final (ca. 3,200-2,400 ya) (Habu, 2004). Thus, the Jomon period spanned over 10,000 years. During the long Jomon period, how did the population sizes change over time? In addressing this question, Koyama (1978) estimated population sizes based on the number of archaeological sites per region and on documentary evidence from the historical period. The results suggest gradual population growth across the Japanese archipelago from the Initial to the Early phase, reaching its peak during the Middle phase, with the eastern part of the archipelago (corresponding to the Kanto region mainly) gaining a larger population size than the western part. However, demographic history during the Jomon period based on archaeological data has remained largely outdated since Koyama’s estimation.

Genetic studies surrounding regional disparities between Eastern and Western Jomon populations have predominantly centered on the prevalence of mitochondrial haplogroups. Among the Jomon people across the Japanese archipelago, three prominent haplogroups—M7a, N9b, and D4b2—have been identified (Shinoda and Kanai, 1999; Adachi *et al*., 2011). Notably, M7a and N9b exhibit low frequencies in the current Japanese archipelago and are virtually absent in present-day populations outside the archipelago. N9b predominates among Eastern Jomon individuals, while M7a prevails among their Western counterparts. The most recent common ancestor (MRCA) of haplogroup M7a is estimated to have existed 30,000 to 20,000 ya, while that of haplogroup N9b is estimated to have lived 23,000 to 5,000 ya. These data suggest a scenario that M7a likely originated from the western regions of the archipelago during the Upper Paleolithic, while N9b may have entered via Northeast Asia, traversing Sakhalin to Hokkaido and the eastern part of the main island (Honshu) during the same period (Shinoda, 2019).

More recently, the Bayesian Skyline plot (BSP) (Drummond *et al*., 2005) analysis has been applied to multiple whole-mitogenome nucleotide sequences from present-day individuals in the Japanese archipelago with the aim of estimating the prehistoric demography (Jinam *et al*., 2021; Mizuno *et al*., 2021). These previous BSPs show a rapid growth in effective populations size after the end of the Jomon period and with the beginning of the Yayoi period, coinciding with the advent of wet rice cultivation. However, since the present-day Japanese gene pool includes mitogenomes from both Jomon people and continental migrants who arrived about 3,000 years ago, BSP analysis of mitogenomes from present-day Japanese does not reflect changes in the population size of the Jomon people.

Population genetic analyses to examine demographic history over thousands of years usually require a minimum sample size of around 50 individuals. However, it has been difficult to achieve this standard using ancient genomes because there are far fewer samples available for analysis compared to modern genomes. Nevertheless, the recent accumulation of complete mitogenome sequences from Jomon bones has opened the way for the estimation of population demography using a relatively large number of Jomon individuals. In this study, we investigated the demographic history of the Eastern and Western Jomon using their complete mitogenome sequence data. We extracted DNA from 53 human remains excavated from Jomon sites and determined 13 new complete mitogenome sequences by high-throughput sequencing (what we call, next-generation sequencing, NGS). Combining previously published mitogenome sequence data, we made a 40-individual dataset, which include 14 Western and 26 Eastern Jomon individuals. We constructed phylogenetic network, estimated diversity statistics, and conducted Bayesian-based demographic analysis using Skyline plots. The results show an increase in effective population size between the Incipient and Initial phases of Jomon period that is not observed in contemporaneous continental East Asian peoples. Notably, this population composition was more pronounced in the eastern part of the Japanese archipelago than in the western part, if we assume that there was no gene flow between East and West, suggesting that the Jomon population exhibited significant regional demographic diversity.

## Material and Methods

### Archaeological samples

53 pieces of skeletal remains excavated from three archaeological sites, Gionbara (GB), Kikumatenaga (KT), and Saihiro (SH) shell mounds, located in Ichihara City, Chiba Prefecture, were examined in this study (Table S1). GB and SH shell mounds were assigned to the Late – Final phases of Jomon period (ca. 4,400 –2,400 ya) and KT shell mound to the Late phase of Jomon period (ca. 3,200 – 2,400 ya). The results of radioisotope dating are described in (Nakamura *et al*., 2024). Hereafter, the three sites will collectively be denoted as Ichihara Jomon sites.

### Radiocarbon dating

Radiocarbon dating was conducted using Accelerator Mass Spectrometry (AMS) at The University Museum, The University of Tokyo. Calibration of dates was performed with reference to IntCal20 and Marine 20 (Reimer *et al*., 2020; Heaton *et al*., 2020) and calibrated ages were determined using OxCal 4.4 software (Ramsey, 2009). Correction for the marine reservoir effect, which causes apparent radiocarbon dates to appear older due to seafood consumption, was achieved by assessing the marine carbon effect based on δ13C values. The marine contribution was estimated by utilizing mean δ13C values (−22.6‰ for land mammal bones and -10.9‰ for marine fish bones) derived from shell middens in Chiba Prefecture, assuming a 5% margin of error. A regional correction value (ΔR) for Tokyo Bay (−98 ± 37 years) was applied to Marine20 based on the analysis of a shell collected in 1882 AD (534 ± 36 BP) (Yoshida *et al*., 2010). The median of the probability distribution of dating was used as the estimated age.

### DNA extraction

All ancient DNA experiments were performed in the clean rooms exclusively built for ancient DNA analyses and installed in the Department of Biological Sciences, Graduate School of Science, University of Tokyo or in the Department of Anatomy, Kitasato University School of Medicine. All bones and teeth were exposed to ultraviolet (UV) light for 15 minutes on either side to reduce DNA contamination from the sample surfaces. Because a sample (GB2-A1a) was calcified and found to be difficult to extract DNA, any further operations were not performed. Petrous bone was cut by the UV-irradiated diamond disc cutter (SHOFU), and a wedge section of the otic capsule region, including “C-part” in Pinhasi *et al*. (2015), was targeted for sampling. The tooth sample was cut by the UV-irradiated diamond disc cutter and separated into crown and root. The dentin powder was collected by UV-irradiated drill (SHOFU). 5-150 mg of bone pieces or powder was collected for DNA extraction.

DNA was extracted following the methods of previous studies (Gamba *et al*., 2014; Damgaard *et al*., 2015; Gakuhari *et al*., 2020). Bone piece or powder was incubated in a 5 mL DNA LoBind tube (Eppendorf) with 2 mL of lysis buffer (Tris HCL pH 7.4 20mM; Sarkosyl NL 0.7%; EDTA pH 8.0 47.5mM; Proteinase K 0.65U/mL) for 15 min at 50 °C at 900 rpm in a Thermomixer (Eppendorf). The samples were centrifuged at 2,300 g for 10 min, and the supernatant was discarded. 2 mL of fresh lysis buffer was added to the tubes and incubated overnight (>16 h) at 60 °C at 900 rpm. After digestion, the tubes were centrifuged at 2,300 g for 10 min. About 2 mL of supernatants were then transferred to ultrafiltration tubes (Amicon® Ultra-4 Centrifugal Filter Unit 10K, Merck). 2 mL of TE buffer was added, and the samples were centrifuged at 2,300 g until the final concentrations reached 100 uL. These concentrates were then transferred to a silica column (MiniElute PCR Purification Kit, QIAGEN) and purified according to the manufacturer’s instructions, except for the elution step with 60 uL EBT buffer (EB buffer with 0.05% TWEEN 20 at the final concentration) pre-heated to 60 °C. The concentration of the DNA extract was measured by Qubit 4 Fluorometer (Invitrogen). The DNA fragment length distribution was measured by Bioanalyzer (Agilent) or TapeStation (Agilent).

### Library preparation

Double-stranded DNA libraries were prepared with NEBNext Ultra II DNA Library Prep Kit for Illumina and NEBNext Multiplex Oligos for Illumina (96 Unique Dual Index Primer Pairs) (New England Biolab: NEB). Around 1 ng of DNA was used for each library. Libraries were constructed according to the manufacturer’s protocol, except for the following two points: the adapter was diluted 10-fold, and size selection was performed. In the size selection, 90 μL of Agencourt AMPure XP (Beckman Coulter) was added to the reaction, and beads with long fragments (>150 bp) were removed by transferring the supernatant to the new 1.5 mL tube. Another 90 uL of AMPure XP solution was added to the supernatant then the short fragments were recovered.

For four samples (KT2, KT68, SH7-3, and SH7-4), Uracil-DNA Glycosylase (UDG) treated libraries (Rohland *et al*., 2015) were prepared. 5 ng of DNA was added to the treatment solution (10X Tango buffer 5 μL; 2.5 mM dNTP 0.2 μL; 100 mM ATP 0.5 μL; 10 U/μL T4 polynucleotide kinase 2.5 μL; 1,000 U/μL USER Enzyme 3 μL; Ultrapure water up to 50 uL in total). The reaction solution was incubated in a thermal cycler at 37°C for 3 h. 1 μL of T4 polymerase was added and incubated at 25°C for 15 min followed by 12°C for 5 min. The product was purified using the Mini Elute PCR purification kit (Qiagen) and extracted twice with EBT buffer warmed to 60°C (first extraction: 30 μL, second extraction: 23 μL), then finally about 50 uL of UDG-treated DNA was obtained. 25 uL of UDG-treated DNA was used for library preparation.

### High-throughput sequencing

Small-scale sequencing was performed on MiSeq (Illumina) to assess the ratios of endogenous human DNA and postmortem damage patterns. We mapped the reads to the human reference genome sequence (hg19) and calculated a mapping ratio (MR), defined as the number of mapped reads divided by the total number of reads. 13 individuals exhibited MR exceeding 15% (refer to Table S1) were subsequently sequenced on HiSeq or NovaSeq (Illumina) with 150 bp or 100 bp paired-end at the National Institute of Genetics, AZENDA, or MACROGEN.

### Raw data processing

The FASTQ files were processed using the following pipeline. For paired-end sequence reads, we used AdapterRemoval v2.3.2 (v.2.2.2 for screening) (Schubert *et al*., 2016) to remove ambiguous and low-quality bases at ends of reads (--trimns and -- trimqualities), and short reads (--minlength 35), combining them into consensus sequence (-collapse). For single-end sequence reads, we used cutadapt v4.4 (Martin, 2011) with options (--trim-n, --quality 2, --length 35) corresponding to those of AdapterRemoval. Using BWA v0.7.17 (Li and Durbin, 2009), the trimmed reads were mapped to the revised Cambridge Reference Sequence (rCRS). In order to map the reads on the end of the reference, we used circularmapper v1.93.5 (Peltzer *et al*., 2016). BAM files from the same library were merged into a BAM file using SAMtools v1.6 (v1.10 for screening) (Li *et al*., 2009). CleanSam and MarkDuplicates in Picard Tools v3.0.0 (v2.21.8 for screening) (https://broadinstitute.github.io/picard/) were used to soft-clip beyond-end-of-reference alignments and remove duplicate reads, respectively. The mapping ratio was calculated using flagstat in SAMtools. The misincorporation patterns were checked by mapDamage2 v2.2.0 (v2.0 for screening) (Jónsson *et al*., 2013) and hard-clipped two or ten bases of 5’ end and ten bases of 3’ end, where substitutions occur frequently, using TrimBam v1.0.15 in BamUtil (Jun *et al*., 2015). For UDG-treated samples, two bases of 3’ and 5’ end of reads were hard-clipped. After merging BAM files from the same individual into a BAM file, mapped reads with mapping quality below Phred score 30 were removed using SAMtools v1.6. We calculated the average depth using DepthOfCoverage in Genome Analysis Toolkit (GATK) v4.3 (Mckenna *et al*., 2010). VCF files were obtained using HaplotypeCaller in GATK and converted to the complete mitogenome sequences in FASTA format using FastaAltenateReferenceMaker in GATK. We applied the same processing pipeline for published ancient genomes in FASTQ or BAM format. The contamination rate was estimated by MitoSuite v1.0.9 (Ishiya and Ueda, 2017). The haplogroup was called by HaploGrep 3 (Schönherr *et al*., 2023).

### Previously published data

27 published whole mitogenome sequences of Jomon individuals (11 Initial, 6 Early, 3 Middle, 5 Late, 2 Final) were used (Fig. 1, Table 1) for comparative analyses. These samples were excavated from archaeological sites across three regions of the Japanese archipelago: 2 from Hokkaido, 24 from Hondo (Honshu, Shikoku, and Kyushu), and 1 from Okinawa. Including our newly sequenced Late Jomon individuals, we obtained a total of 40 Jomon individuals’ whole mitogenome sequences (11 Initial, 6 Early, 3 Middle, 18 Late, 2 Final) (Table 1). The Jomon individuals were assigned to Eastern or Western Jomon depending on whether the site where they were excavated was located on the east or west of the Fossa Magna, which is the central rift valley of the Japanese archipelago (Fig.1). One Upper Paleolithic individual from Okinawa (Minato 1) and one modern individual from Sudan in Northeast Africa (JN655840.1) were used in the phylogenetic analysis (Table 1).

**Fig. 1.**
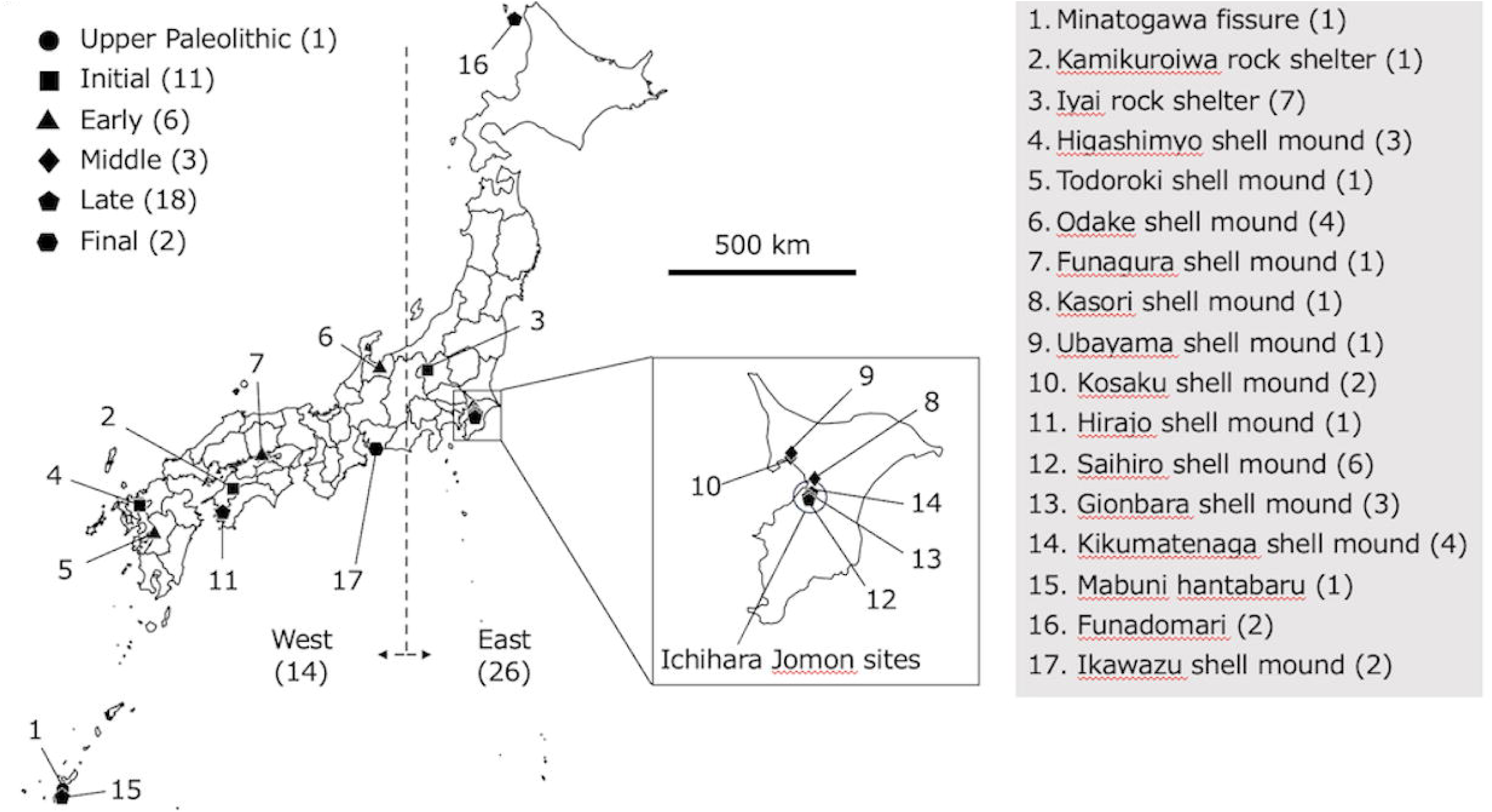
Archaeological sites from which the ancient samples were examined in the study. Each site is marked with a circle, square, triangle, rhombus, pentagon, and hexagon, representing the Upper Paleolithic period, Initial, Early, Middle, Late, Final phases of the Jomon period, respectively. The square shows an enlarged view of Chiba Prefecture that the 13 individuals we newly sequenced were excavated. We defined East and West of the Japanese archipelago bordered by Fossa Magna.

**Table 1.**
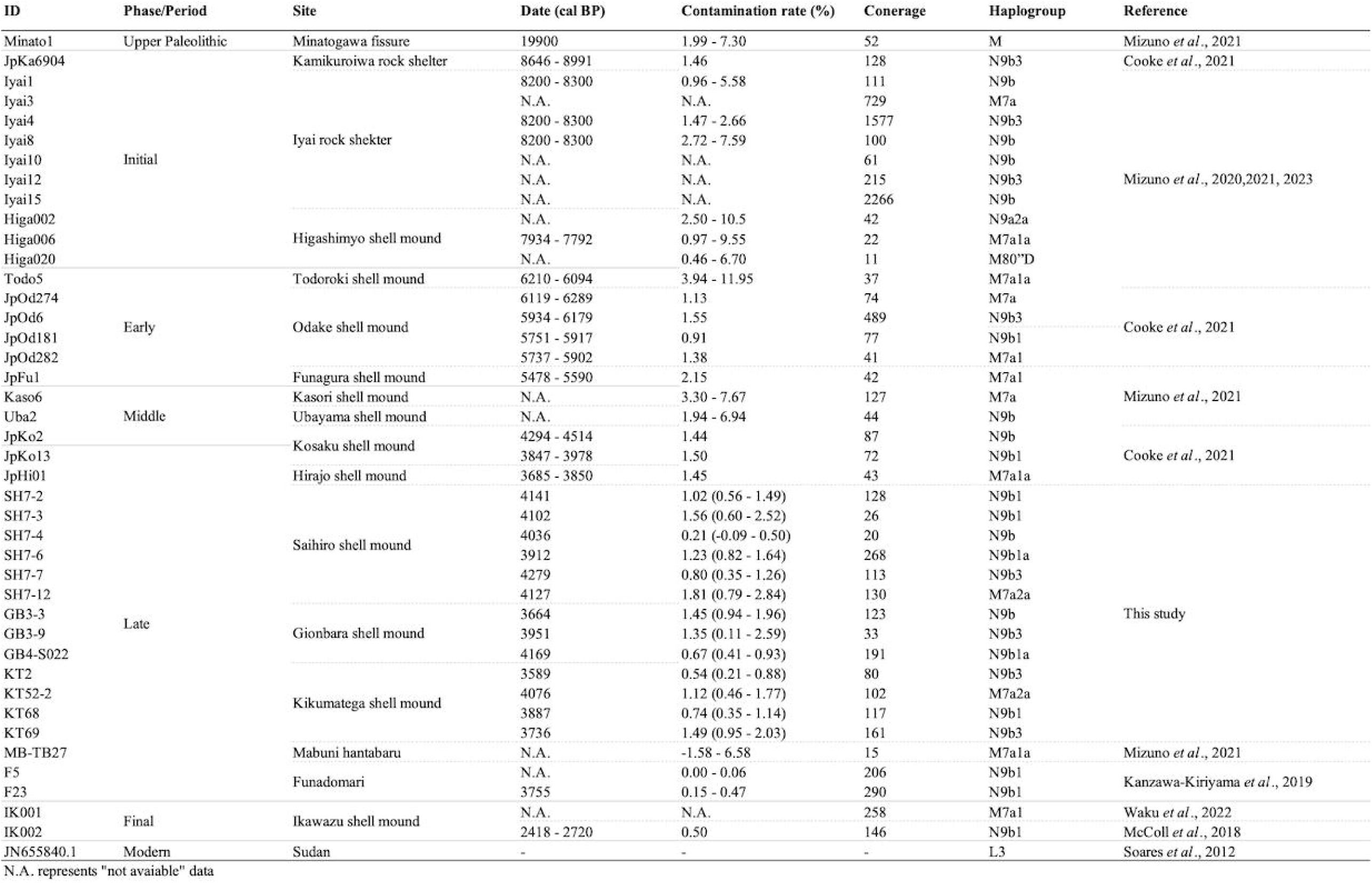
List of ancient and modern human samples.

We aligned all FASTA-formatted sequences to rCRS using MEGA X (Kumar *et al*., 2018). We used hyper-variable (HVI and II) regions that correspond to the nucleotide position of 1-576 and 16024-16569 of the rCRS, respectively, as neutral regions. Because of difficulty in alignment (owing to repetitiveness and/or indel), we removed the nucleotide positions, 303-315, 522-523, 16180-16193, and 16519 of rCRS from the sequence data set for the subsequent analyses.

### Phylogenetic analysis

A median-joining network (Bandelt *et al*., 1999) was created based on nucleotide sequences of whole-mitogenome using POPART ver. 1.7 (Leigh *et al*., 2015). We added an Upper Paleolithic and a present-day individual as an outgroup to root the tree (Table 1).

### Genetic diversity statistics

Using DnaSP ver. 6.12.03 (Rozas *et al*., 2017), we calculated the values of genetic diversity statistics, nucleotide diversity and Tajima’s *D*. Since these statistics assume the neutral theory, we used hyper-variable regions (HVI and II, 1094 bp) for calculation. Nucleotide diversity is the mean pairwise difference of nucleotides divided by the sequence length and indicates the genetic diversity within a population (Nei, 1987). Tajima’s *D* evaluates the difference between nucleotide diversity and the proportion of polymorphic sites adjusted for the sample size (Tajima, 1989). Although Tajima’s *D* was originally developed for testing the neutral hypothesis in molecular evolution, it is also used to detect change of population size because Tajima’s *D* is based on the expectation of a constant population size at mutation drift equilibrium (Aris-Brosou and Excoffier, 1996; Oota *et al*., 2002a; Mousset *et al*., 2004; Matsukusa *et al*., 2010; Katsumura *et al*., 2012; Puzey *et al*., 2017; Flanagan *et al*., 2021; Peng *et al*., 2023). Using the HVI and II regions of the mitogenome, nucleotide diversity and Tajima’s D were calculated for the 40 Jomon individuals or divided into groups, as well as for 103 CHB individuals (Chinese from Beijing) and 104 JPT individuals (Japanese from Tokyo) in phase 3 (1000 Genomes Project Consortium, 2015).

### Bayesian skyline plot to estimate demographic history

Coalescent theory provides a framework for understanding the relationship between demographic history of a population and genealogy (Pybus *et al*., 2000). In human population genetics, effective population size and demography were initially estimated based on pairwise mismatch distribution by comparing haploid sequences such as the mitogenome and Y chromosome within populations (Rienzo and Wilson, 1991; Harpending *et al*., 1998). In more theoretical studies, a method for estimating demography from phylogenetic information was proposed (Fu, 1994), and a coalescent model was introduced to estimate the variable size of past populations from modern nucleotide sequence data (Griffiths and Tavare, 1994; Donnelly and Tavare, 1995). The lineages-through-time (LTT) plot is a method of displaying the proportion of congeneric to phylogenetic trees (Nee *et al*., 1995), and the Skyline plot, one of LTT plots, is a population-size piecewise constant model that can be adapted to a wide range of demographic scenarios (Pybus *et al*., 2000). It is useful as a model selection tool to indicate the most appropriate demographic model for any given data set (Pybus *et al*., 2002).

To investigate the change in effective population size of the Jomon population, we employed a Bayesian skyline plot (BSP) using BEAST ver2.7.5 (Bouckaert *et al*., 2019). BSP is based on Bayesian coalescent inference of multiple haploid genomes experiencing uniparental inheritance without recombination. The input XML files for BEAST were created by BEAUTi program, given the tip date (Table 1). For individuals for which calibration dates were unavailable, we referred to the sub-period to which they belonged or the date of other individuals from the same site. We used a gamma-distributed rate to apply the general time reversible model mutation model. Following a previous study, we set a clock model to a log-normal distribution and molecular clock to 2.77 × 10^−8^ (Posth *et al*., 2016). Markov Chain Monte Carlo (MCMC) was set with the chain length of 1 × 10^7^ with 1 × 10^5^ burn-in steps to collect sufficient samples for parameter estimation. After the analyses by BEAST, we put the output log files to Tracer ver. 1.7.2 (http://tree.bio.ed.ac.uk/software/tracer/) to draw BSP.

## Results

### Data quality of the newly sequenced individuals

We obtained DNA from 52 pieces of skeletal material at the Ichihara Jomon sites. The mapping rate (MR), the rate of the reads to the human reference genome out of raw sequence reads, was between 0 and 62.97%, and 13 individuals exceeded 15% (refer to Table S1). We further sequenced these 13 individuals and obtained complete mitogenome sequences with average depths of 20–270-fold and contamination rates below 2% (Table 1). Filtered reads showed deamination at the reads ends (see Figs. S2).

### Phylogenetic network and major haplogroups distribution

To elucidate the phylogenetic relationships among 40 Jomon individuals (Table 1 and Fig.1), we constructed a phylogenetic network based on whole-mitogenome sequences (Fig. 2). The network showed two prominent clades corresponding to haplogroup N9b and M7a, encompassing 26 and 12 individuals, respectively. Consistent with findings of a previous study (Mizuno *et al*.,2023), only two (Higa002 and 020 from the Initial phase) out of the 40 Jomon individuals, along with the Upper Paleolithic Minatogawa individual (Minato1), did not fall into the two major haplogroups.

**Fig. 2.**
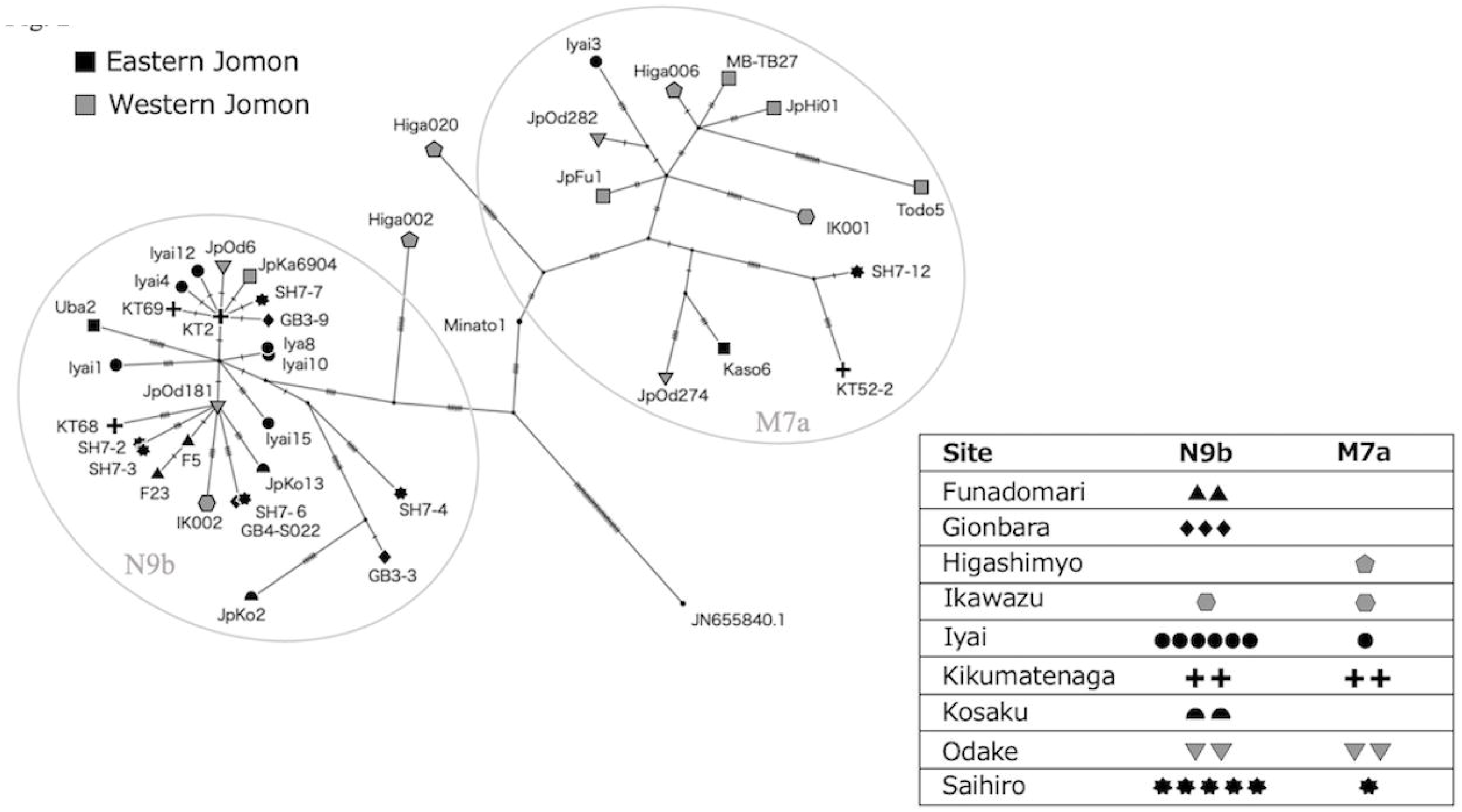
The phylogenetic network of whole-mitogenome sequences using the median-joining method. Eastern Jomon individuals are colored in black, and Western are in gray. Each mark except for Square represents an archaeological site that includes multiple individuals, whereas Squares represent a site that include only a single individual. Among the 40 mitogenome sequences, the pairs of sequences from six individuals were identical to each other. The three pairs are described as SH7-2/SH7-3, SH7-6/GB4-S002, and Iyai8/Iyai10. The number of hatch marks on each branch is that of mutation. The box on the right shows the number of individuals belonging to haplogroups N9b or M7a at sites where multiple individuals were used in this study. There is a single individual from Higashimyo in M7a, because Higa002 and Higa020 did not belong to either N9b or M7a, although three individuals from Higashimyo were examined in this study.

Previous studies on the Jomon mitogenome haplogroups suggested that a higher prevalence of N9b in the eastern region of the archipelago and M7a in the western region (Adachi *et al*., 2008; 2013; 2021; Mizuno *et al*., 2023). To verify this east-west dichotomy in haplogroup distribution using the whole-mitogenome sequences, we categorized the 31 individuals from nine sites into either N9b or M7a (Fig. 2). Four sites detected only either N9b or M7a, while five sites detected both haplogroups. An analysis of the frequency distribution revealed that 22 out of 26 Eastern Jomon mitogenomes belonged to N9b, whereas 8 out of 12 Western Jomon mitogenomes belonged to M7a. We performed a Fisher’s exact test for 38 individuals who belong to N9b or M7a, and detected a significant association of N9b with Eastern Jomon individuals and M7a with Western Jomon individuals (p<0.005). Thus, while these haplotypes were not strictly confined to specific regions, there existed statistically significant disparities in the frequency distribution of N9b and M7a between the eastern and western populations.

Examining the phylogenetic network’s topology, we observed star-like structures, where multiple sequences diverged from a single sequence, within the N9b clade but not the M7a clade. The presence of such star-like structures typically indicates rapid population expansion (Rienzo and Wilson, 1991; Oota *et al*., 2002b). This suggests that a rapid demographic expansion occurred within the lineage of N9b or within the Eastern Jomon population (as opposed to the Western population).

### Diversity statistics to test demographic expansion

To elucidate the likely demographic expansion within the Jomon population, we computed diversity statistics based on nucleotide sequences from the HVI region of the mitogenome. Table 2 presents the number of segregation sites (S), population mutation rate (Θ), nucleotide diversity (π), and Tajima’s *D* for the groups categorized by archaeological site(s) (Iyai, Higashimyo, Odake, and KT GB, SH, combined as Ichihara) and by geographical region(s), comparing them with modern East Asian populations (CHB and JPT). Among the groups, excluding the modern populations, the minimum π value was 0.00279 (Iyai), while the maximum was 0.00439 (Higashimyo). Both of the values were from the Initial phase of the Jomon period. Eastern and Western Jomon exhibited nearly identical π values (0.00310 and 0.00314, respectively), indicating comparable genetic diversity between the eastern and western regions of the archipelago. Notably, the π value of all Jomon individuals (0.00330) was considerably lower than that of CHB and JPT (0.00884 and 0.00785, respectively), suggesting a relatively reduced genetic diversity among the Jomon people compared to modern populations.

**Table 2.**
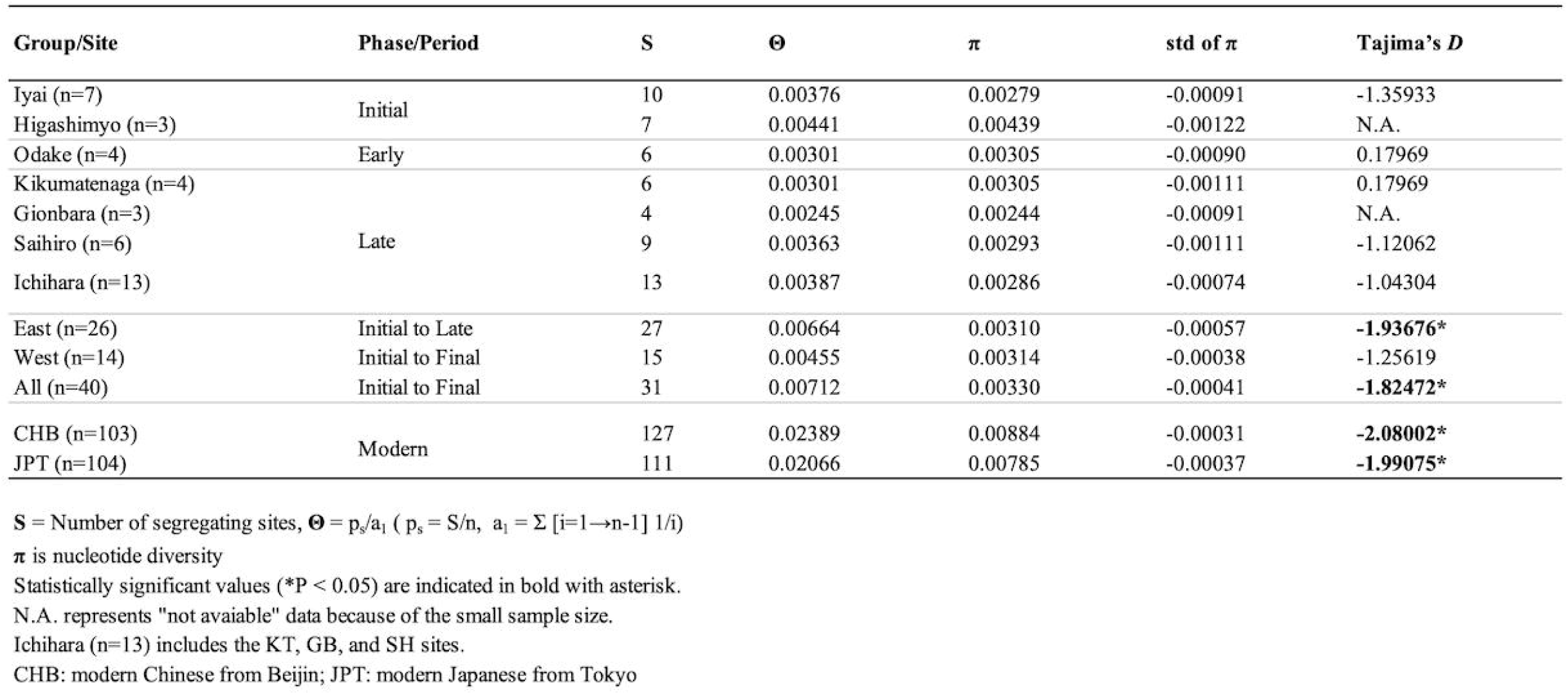
Statistical summary of the Jomon groups.

Tajima’s *D* test was performed assuming neutrality for the HVI and II regions of the mitogenome (Table 2). Tajima’s *D* describes the relationship between the number of segregation sites (S: the proportion of polymorphic sites) and nucleotide diversity (π) under the equilibrium of genetic drift and mutation. Under the neutrality, the ratio of S to the number of samples is equal to π. A Tajima’s *D* value greater than zero suggests a higher proportion of sites with elevated nucleotide diversity compared to neutrality, while a negative value suggests a greater proportion of sites with reduced nucleotide diversity. When Tajima’s *D* significantly deviated from zero in a negative direction, it indicates a rapid population expansion, while non-significance suggests a constant population size. Both CHB and JPT exhibited significantly negative value, indicative of demographic expansion in modern populations. Each Jomon group showed no significant deviation from zero, suggesting a constant population size within each local community. This result did not change even when considering Western Jomon as a collective group. However, when aggregating Eastern Jomon and all Jomon individuals from each site, the Tajima’s *D* values were significantly negative. Thus, it is likely that overall, the Jomon population experienced a substantial expansion, particularly evident in the Eastern Jomon population.

### Demographic change in the Jomon period

To assess over-time changes in the effective population size (*N*e), we conducted the BSP analyses using whole-mitogenome sequences (Fig. 3). In the BSP encompassing all Jomon individuals, we found that a pronounced increase in *N*e occurred approximately 13,000 – 8,000 ya, matching the transition from the Incipient to the Initial phase of the Jomon period (Fig. 3A). Assuming no gene flow between East and West, we conducted BSPs with dividing Eastern and Western Jomon individuals. Then we found the demographic growth of Eastern Jomon was particularly remarkable, while that of Western Jomon was comparatively moderate (Fig. 3B). Interestingly, such demographic growth during the Incipient and Initial phase was absent in the BSP of the present-day Japanese individuals, which instead indicated a demographic growth around 5,000 – 2,000 ya, corresponding to the Middle to the Final phases of the Jomon period (Fig. 3C). This discrepancy likely arises from the present-day Japanese gene pool incorporating mitogenomes from both Jomon people and continental populations who migrated around 3,000 ya. Consequently, we infer that demographic growth occurred during the transitional period from the Incipient and Initial phase among the Jomon people but not among continental East Asian populations.

**Fig. 3.**
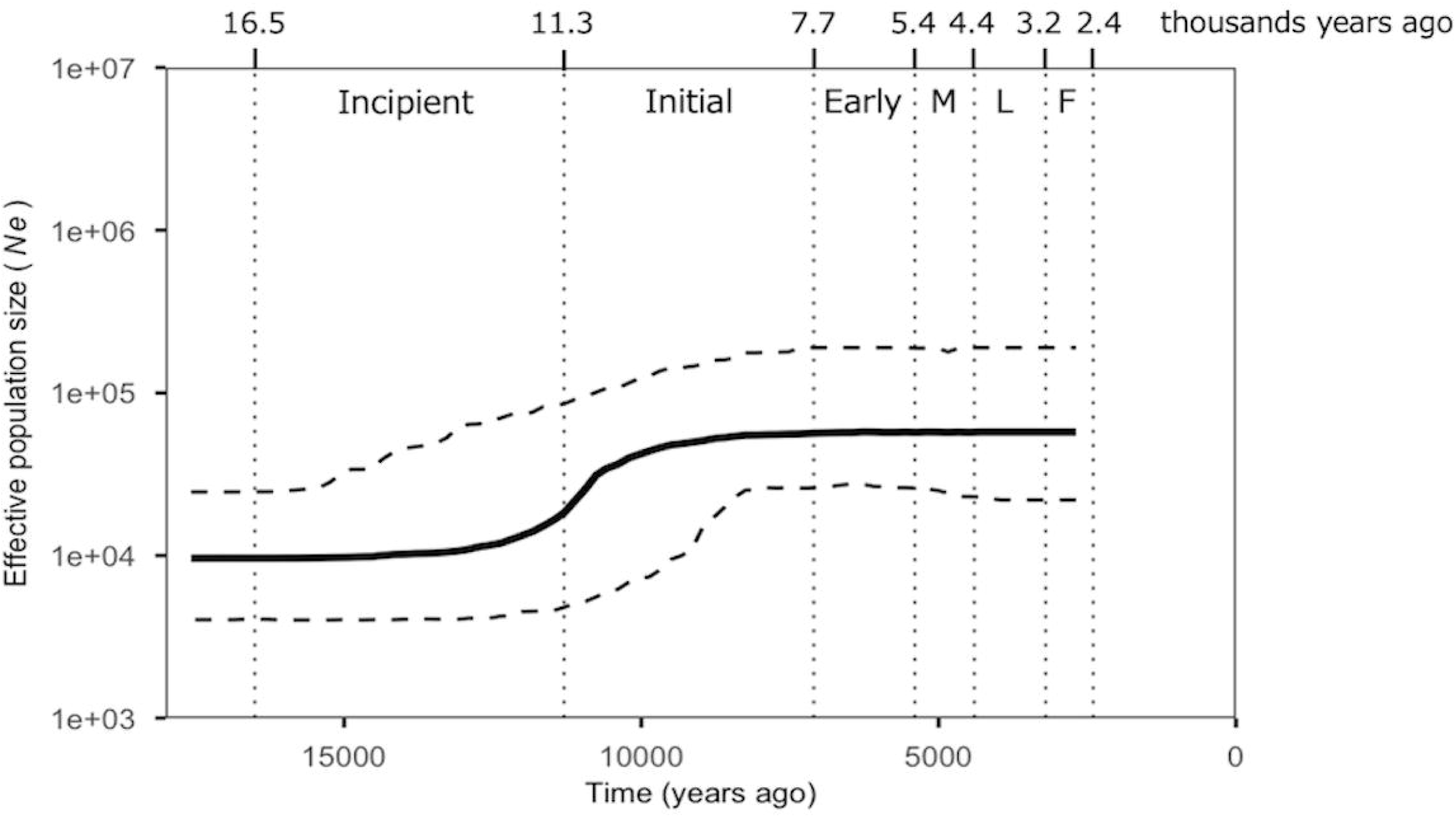
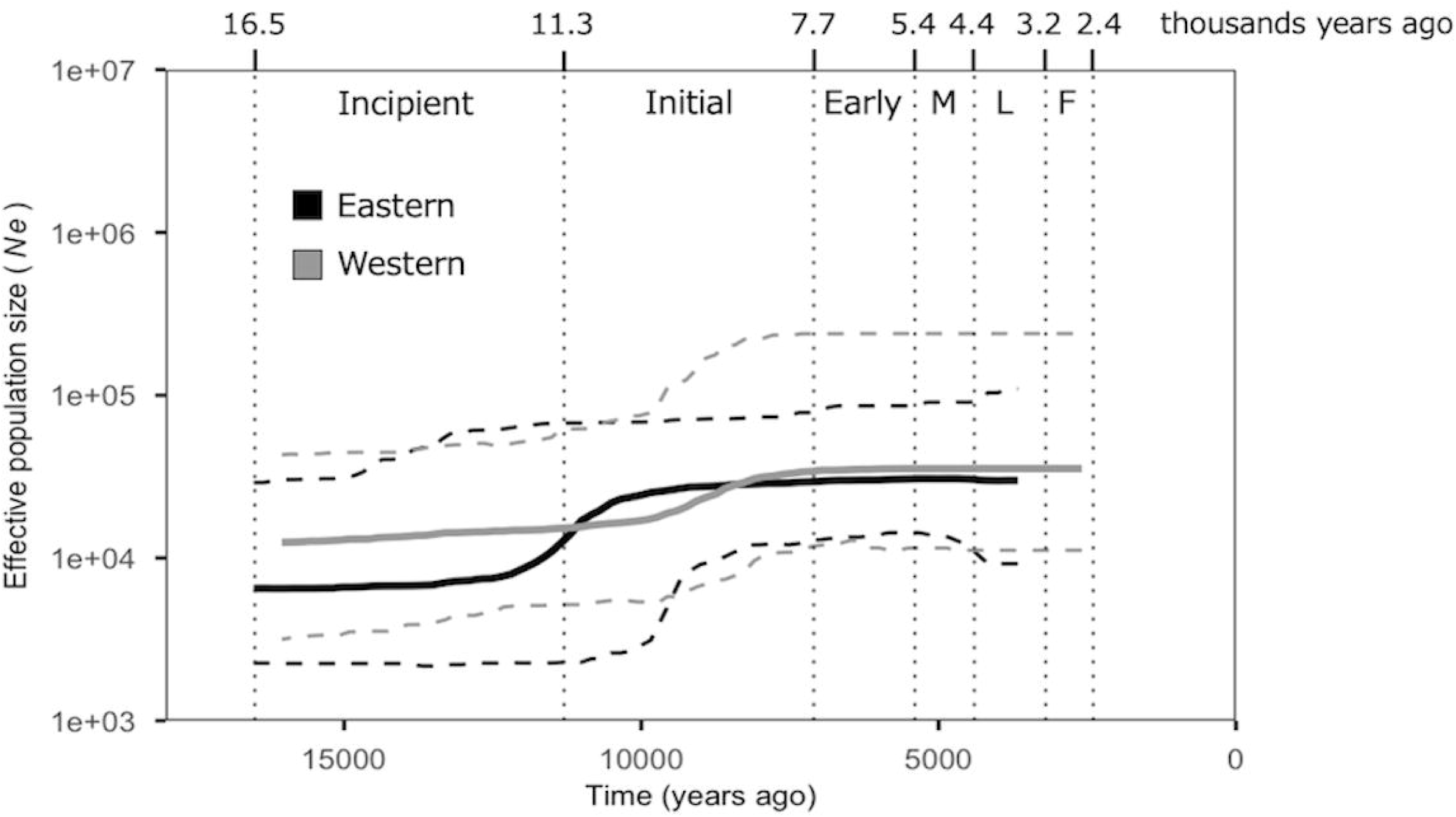
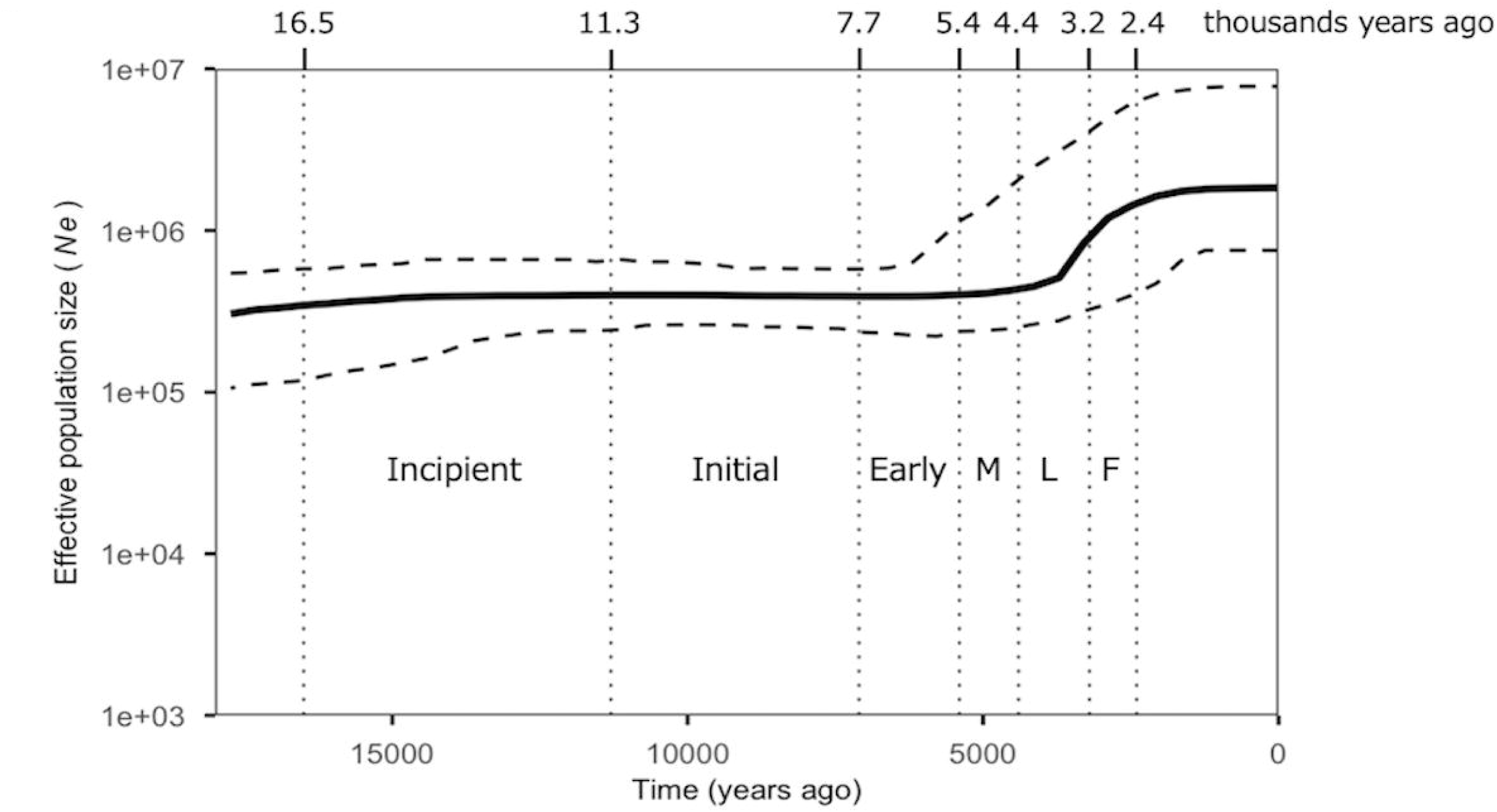
Bayesian Skyline Plot of all Jomon (A), Eastern (black) and Western (gray) Jomon individuals (B), and JPT (C). Time on the x-axis is shown in years ago, and effective population size on the y-axis is shown on the log scale. The bold line indicates the estimated median effective population size, the two dashed lines across it indicate 95% credible intervals, and the dotted line parallel to the y-axis indicates the boundary between the six phases, Incipient, Initial, Early, Middle(M), Late(L), Final(F) of the Jomon period.

## Discussion

In this study, we delved into the demographic history of the Jomon people through comprehensive analysis of whole-mitogenome sequence data, involving the Initial, Early, Middle, Late, Final phases of the Jomon period, from the Hokkaido, Hondo, and Okinawa in the Japanese archipelago (Fig.1 and Fig. S1). While nuclear genome sequence data from ancient specimens has become more accessible, thus offering avenues for exploring the demographic history of the Jomon people, it is important to notice the inherent differences between nuclear and mitochondrial genomes. Each mutation (SNP: single nucleotide polymorphism) in the nuclear genome possesses a coalescent time older than that in the mitogenome, owing to the difference of the effective population sizes between diploid and haploid genomes. Consequently, demographic changes estimated based on nuclear genome data pertain more to the Pleistocene rather than the Holocene. Meanwhile, when we want to look into demography in the Holocene, it is likely that mitogenome data are more informative than nuclear genome data.

The analytical depth presented in this study was made possible by the accumulation of whole-mitogenome sequence data from multiple Jomon individuals over the last several years. However, analyzing Jomon demography using the whole-mitogenome sequence data of modern individuals residing in the Japanese archipelago remains challenging due to the substantial presence of mitogenomes from individuals who migrated from the East Asian continent 3,000 years ago. Identifying mitogenomes specific to the Jomon people from the modern Japanese gene pool is difficult (Watanabe *et al*., 2019), thus hindering the clarification of demographic history in the maternal lineage preceding the Jomon period. Notably, a previous study has successfully identified a Y chromosome specific to the Jomon people. Watanabe *et al*. (2019) conducted a phylogenetic analysis of the East Asian Y chromosome, identifying a major clade (comprising 35.4% of Hondo Japanese) exclusively composed of the Japanese Y chromosome. Monte Carlo simulations revealed that approximately 70% of Jomon males possessed Y chromosomes belonging to this clade, with BSP analysis indicating a notable decline in the male population around 2,500 years ago, aligning with the transition from the end of the Jomon to the early Yayoi period.

The present study allowed us to discuss the demography of the Jomon people not only through the paternal lineage but also through the maternal lineage. However, due to limited data from individuals in the Late phase of the Jomon period and absence of individuals from the Yayoi period in this analysis, the pronounced phenomenon of effective population size observed around 2,500 years ago on the Y chromosome could not be corroborated with the mitogenome data. Hence, the ongoing debate regarding the observation of such a phenomenon remains unresolved. In the near future, the addition of whole-mitogenome sequence data from the Late phase, Yayoi, and even Kofun periods holds promise for facilitating comparisons between paternal and maternal lineages.

The phylogenetic network, constructed using whole-mitogenome sequence data, revealed two distinct clades corresponding to M7a and N9b (Fig. 2). Notably, star-like structures were observed exclusively within the N9b clade, suggesting a potential signal of demographic expansion. Consistent with previous studies, when dividing Jomon individuals into Eastern and Western groups using the Fossa Magna as a boundary, the N9b clade predominantly encompassed Eastern Jomon individuals, whereas the M7a clade mainly comprised Western Jomon individuals. Despite this division differing from conventional archaeological delineations between East and West, there were no discernible differences in the plot of Eastern Jomon, even when employing alternative borders such as the Chubu-Kinki boundary suggested by Koyama (1978), while that of Western Jomon was not comparable because of the small sample size (Fig. S3). The population growth patten of the Eastern Jomon with Koyama’s grouping was similar to ours, suggesting that the demographic expansion in the Kanto region accounts for a large portion of that of Jomon people as a whole. However, the fact that our estimate of the timing of population growth in this study is older than Koyama’s estimate is still an open question. It should be noted that the Jomon individuals used in this study are concentrated in the Kanto region (24 of 40 individuals), and thus it cannot be denied that the demographics of this region may be regarded as those of the eastern part of the Japanese archipelago. Further work will be required to collect genome sequences of the Jomon people from the broader area in the archipelago.

To reinforce these findings, we conducted BSP analyses. When considering all 40 Jomon individuals as a single group, the population exhibited a notable increase from the Incipient to the Initial phase (Fig. 3A). Upon division into Eastern and Western groups, Eastern Jomon displayed a distinct population expansion from the Incipient to the Initial phase, while Western Jomon showed a more moderate population growth during the middle of the Initial phase (Fig. 3B). This population growth pattern was not evident in BSP analyses utilizing whole-mitogenome sequences from modern Japanese. Namely, the demographic expansion during this period is thought to have occurred uniquely to the Jomon population, distinct from continental East Asians. This finding carries significant implications for the prevailing scenario regarding the peopling history of the Jomon population based on haplogroups M7a and N9b.

According to this scenario, it has been proposed that at least two ancestral lineages of Jomon people reached the Japanese archipelago: N9b, likely originating from Northeast Asia via Sakhalin and Hokkaido, and M7a, possibly arriving from the West through the Korean peninsula (Shinoda, 2019). The scenario is based on the idea that the disparity in the frequency distribution of M7a and N9b between the Eastern and Western regions reflects such distinct migration routes. However, our results analyzed on whole-mitogenome sequences from 40 Jomon individuals contradict this scenario. The coalescent times estimated from the sequence data involved in these haplogroups were approximately 10,000 years ago (refer to Fig. S4). Considering that the Jomon period commenced roughly 16,000 years ago, it is conceivable that M7a and N9b are haplogroups that originated within the Japanese archipelago. This inference is supported by the fact that among the 40 individuals analyzed in this study, two individuals from the Higashimyo site (Higa002 and Higa020) of the Initial phase and the Minatogawa individual (Minato1) from the Upper Paleolithic period were notably not classified into M7a and N9b but instead located on branches closer to the root (see Fig. 2).

If indeed both haplogroups originated within the Japanese archipelago, the aforementioned scenario would be rejected, and the geographical difference in frequency distribution between the East and West could be attributed to genetic drift, rather than different migration routes. The idea that the difference in the frequency of the two haplogroups in the East and West was caused by genetic drift is based on the assumption that the gene flow between the East and West Jomon was interrupted for a period of time. But does the archaeological evidence support such a situation? Further discussion between archaeology and genetic anthropology is necessary. Previous studies based on modern mtDNA HV region have estimated the most recent common ancestor (MRCA) of these haplogroups to be 20,000 to 10,000 years older (Tanaka *et al*., 2004) than the coalescent times we estimated in this study. The reason for this discrepancy remains unknown. To address it, increasing the dataset of Jomon individuals from the Incipient and Initial phases, as well as from the western region of the archipelago, will be crucial.

Archaeological observations have long noted disparities in site numbers between the East and West, with this gap purportedly widening significantly during the Middle phase (Koyama, 1984). However, the timing of population growth we identified precedes this period by nearly 5,000 years. While age estimates based on genome information contain relatively large margins of error in general, the underlying cause of this discrepancy remains elusive. In archaeological contexts, efforts have been made to correlate the East-West disparity in site numbers, indicative of population size, with regional differences in natural environments and ecosystems. Eastern Japan, characterized by deciduous broad-leaved forests dominated by beech, contrasts with western Japan’s evergreen broad-leaved forests centered around oak. Yamauchi (1969) proposed the hypothesis that salmon and trout served as crucial animal resources in Eastern Japan (Taniguchi, 2019). In the next step of this research field, it will be most important to interpret Jomon genome information based on these ecological perspectives.

## Data Availability

All sequence data determined in this study have been deposited in the DNA DataBank of Japan (DDBJ) (Accession No. LC805243-LC805255)

## Acknowledgments

We thank to the helpful comments from Drs. Adrian Davin, and Steven Abood in University of Tokyo. This study was mainly supported by JSPS KAKENHI Grant-in-Aid for Scientific Research 18H03590 and 22H00020 to R.T. This study was also supported by JSPS KAKENHI Grant-in-Aid for Scientific Research 23H04838, 23KJ0654, 21H04779, 21H00337, 21H05362, 21K19289, 19H04526.

