## Supplementary material for "Endemic demographics of the Jomon people estimated based on complete mitogenomes reveal their regional diversity": Fig. S1-S4

### Slide 1
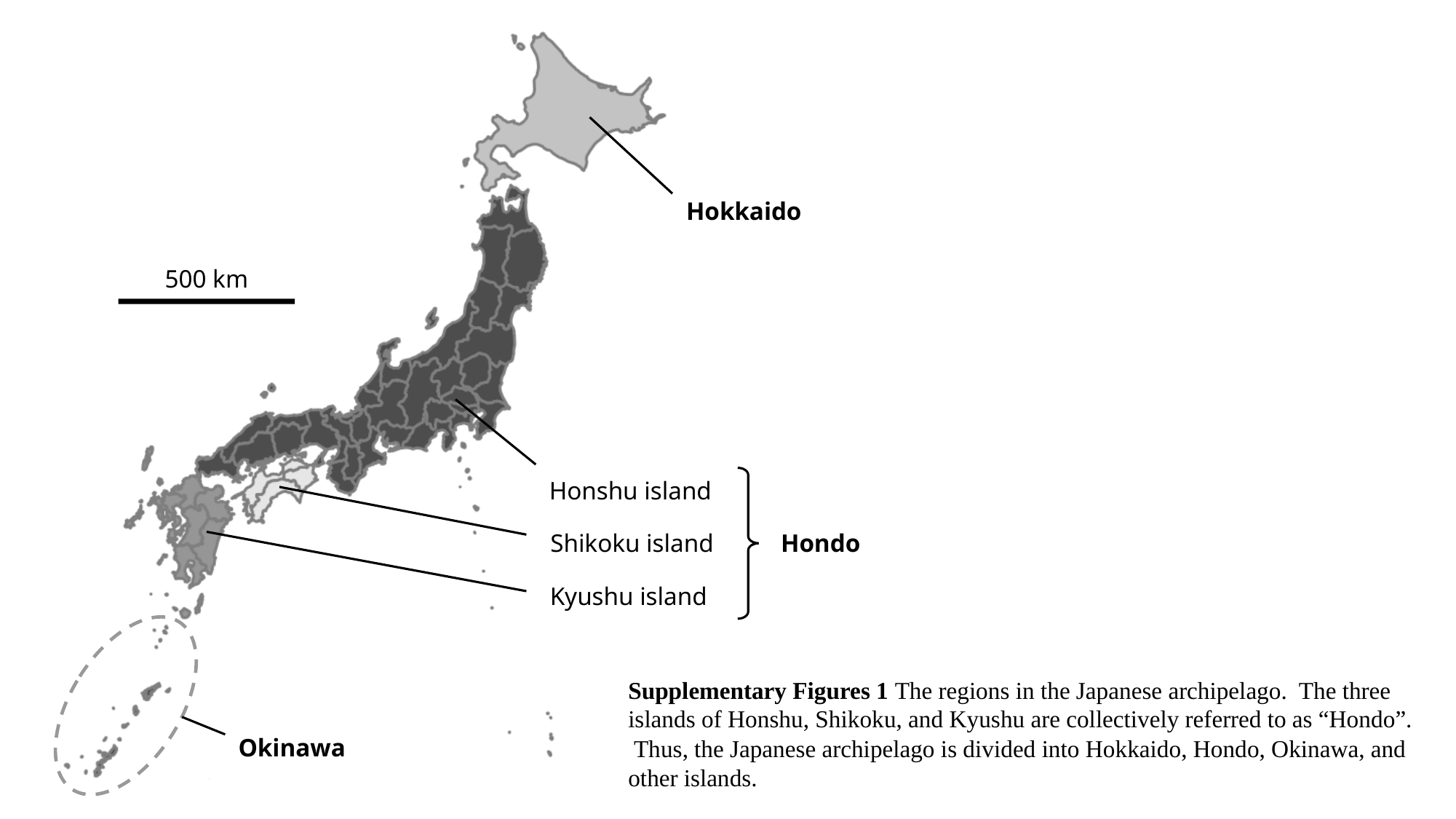

Hokkaido
Honshu island
Shikoku island
Hondo
Kyushu island
Okinawa
500 km
Supplementary Figures 1 The regions in the Japanese archipelago. The three islands of Honshu, Shikoku, and Kyushu are collectively referred to as “Hondo”. Thus, the Japanese archipelago is divided into Hokkaido, Hondo, Okinawa, and other islands.

### Slide 2
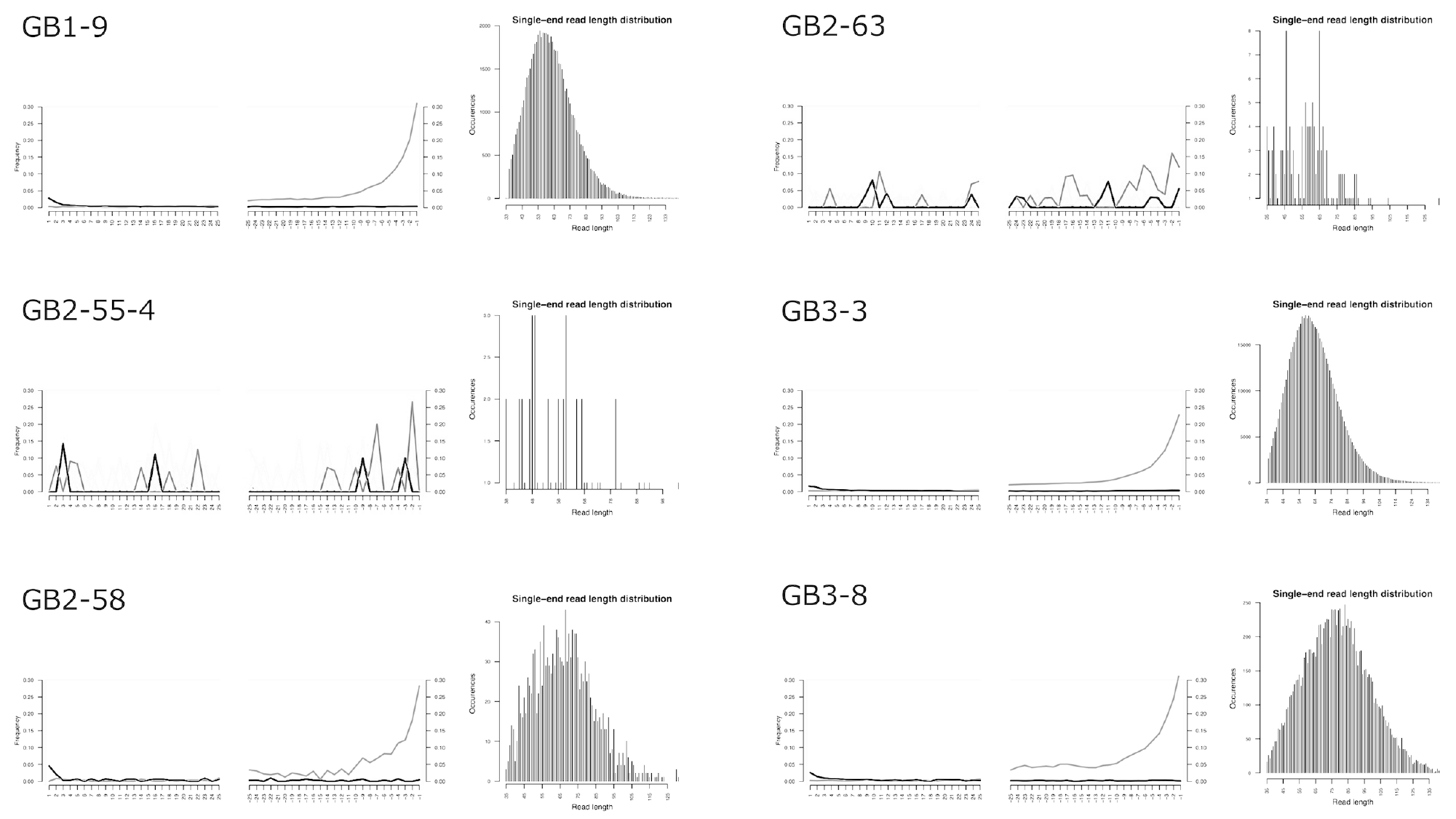

### Slide 3
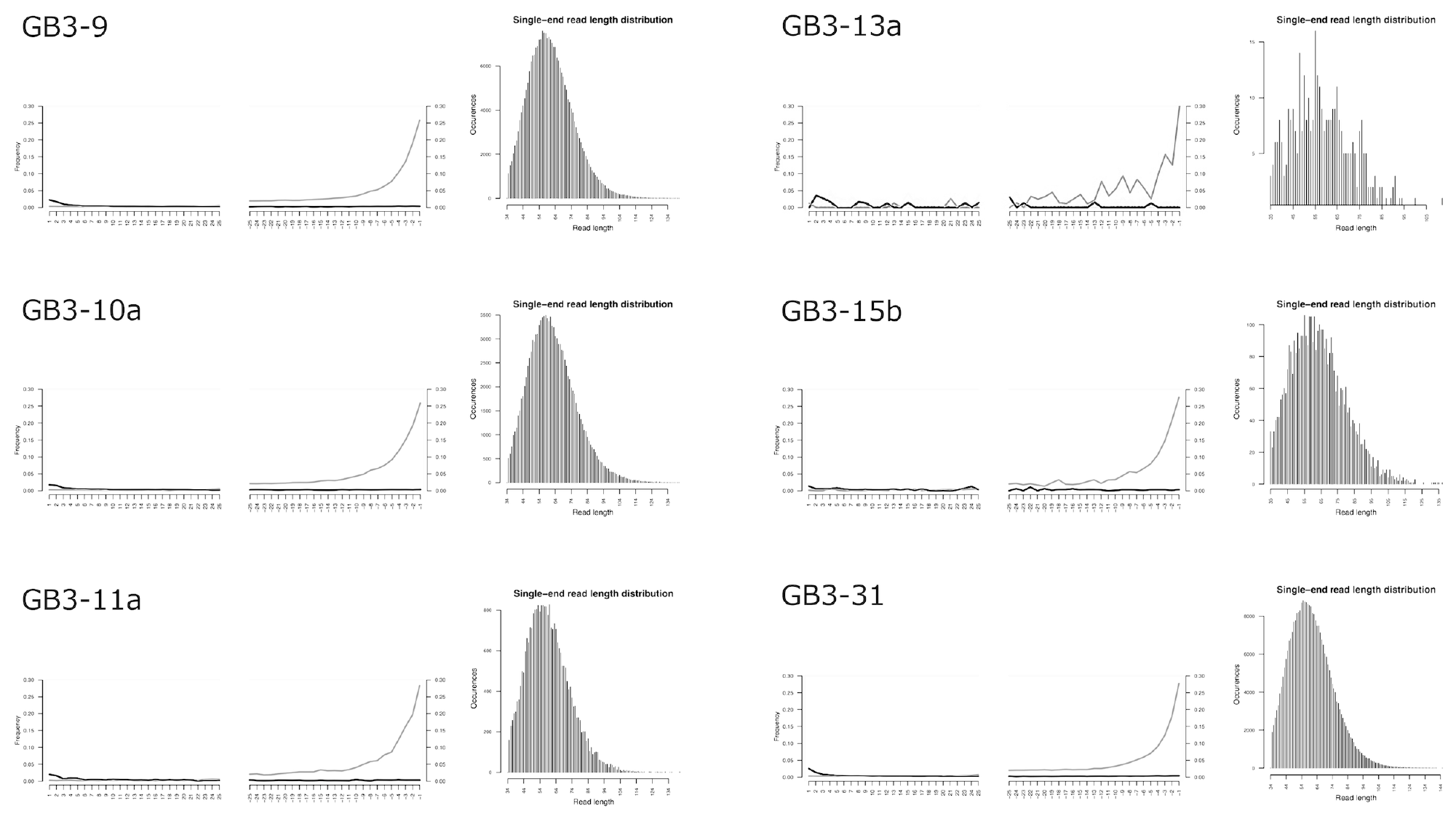

### Slide 4
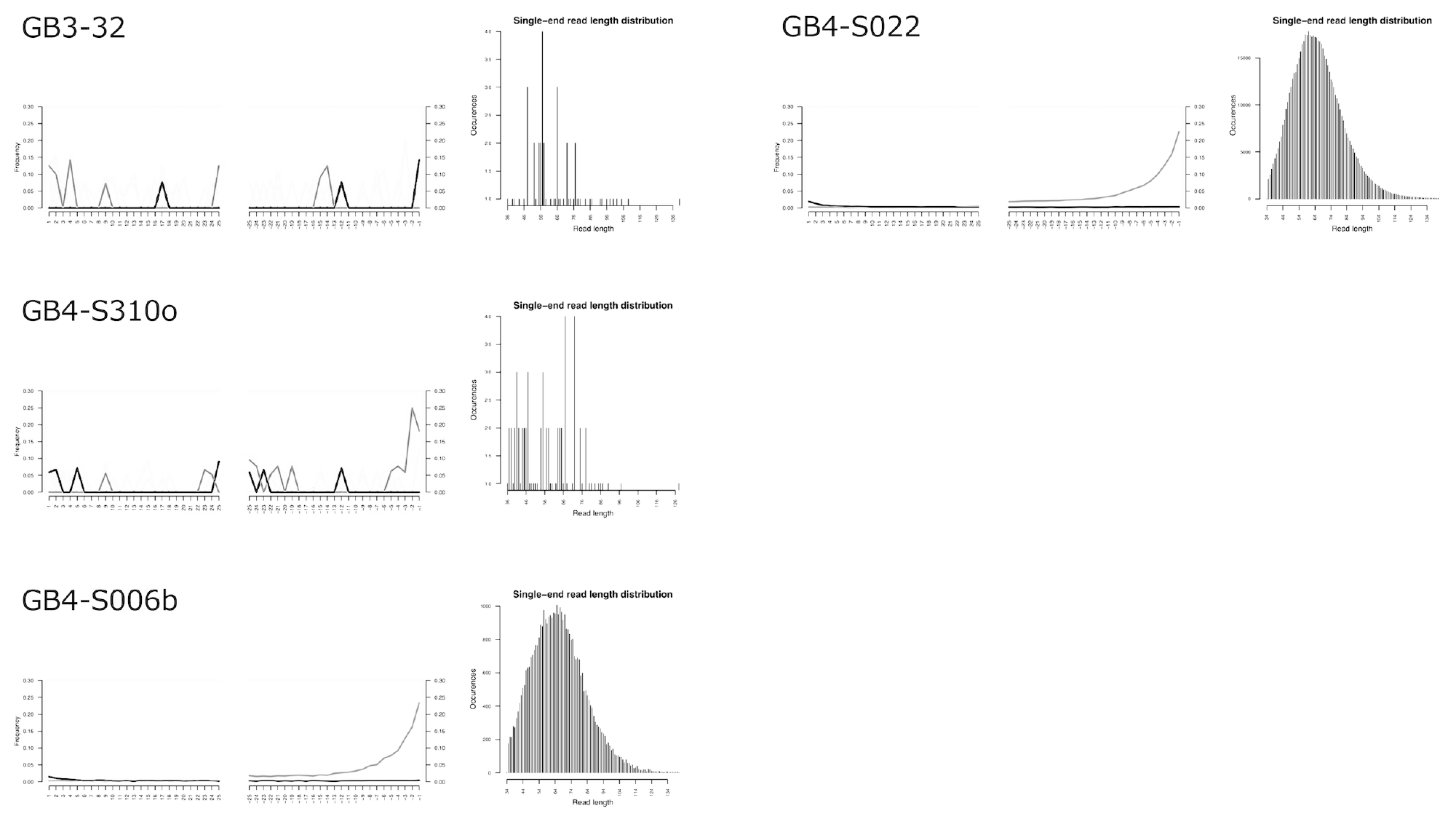

### Slide 5
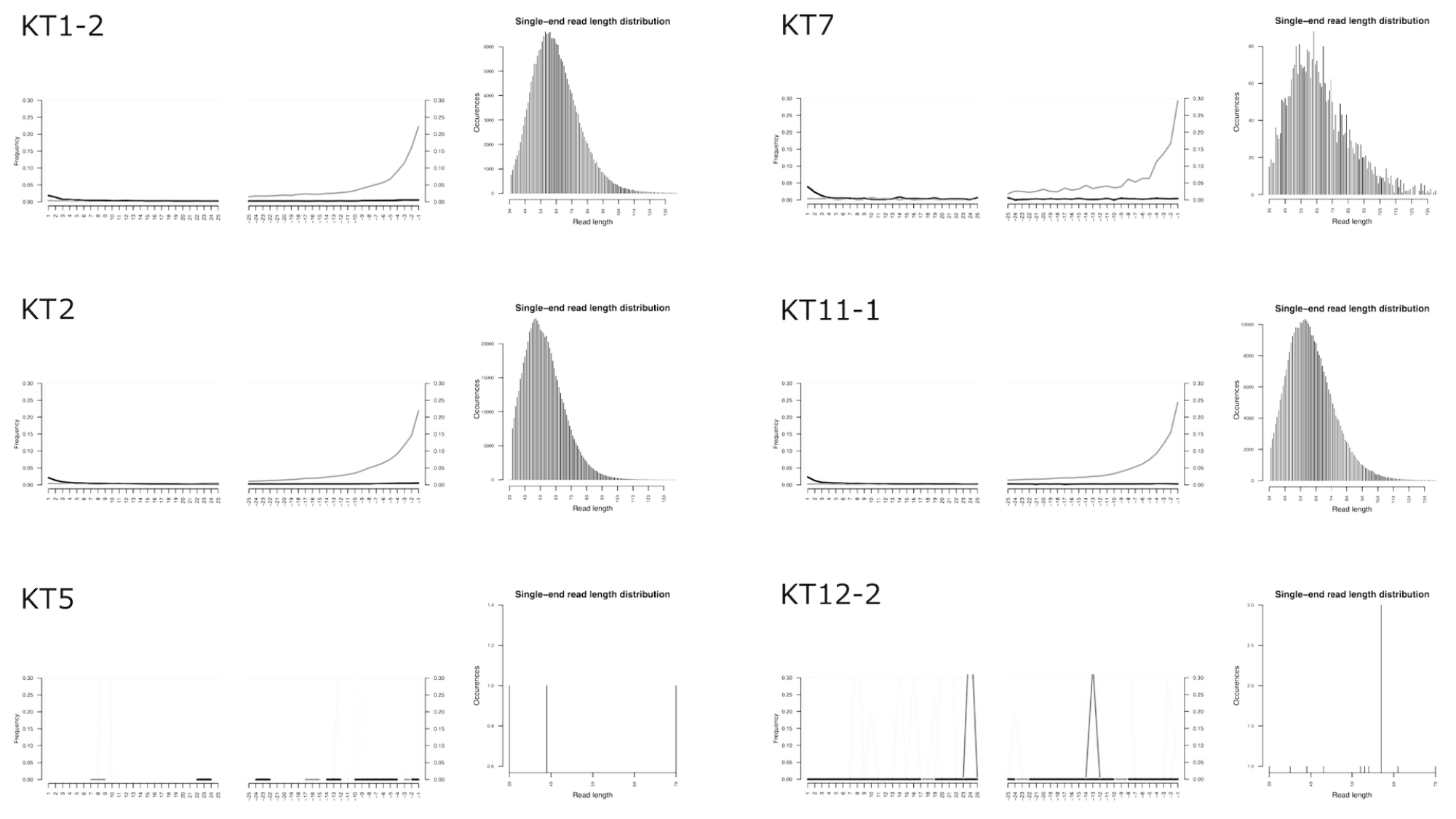

### Slide 6
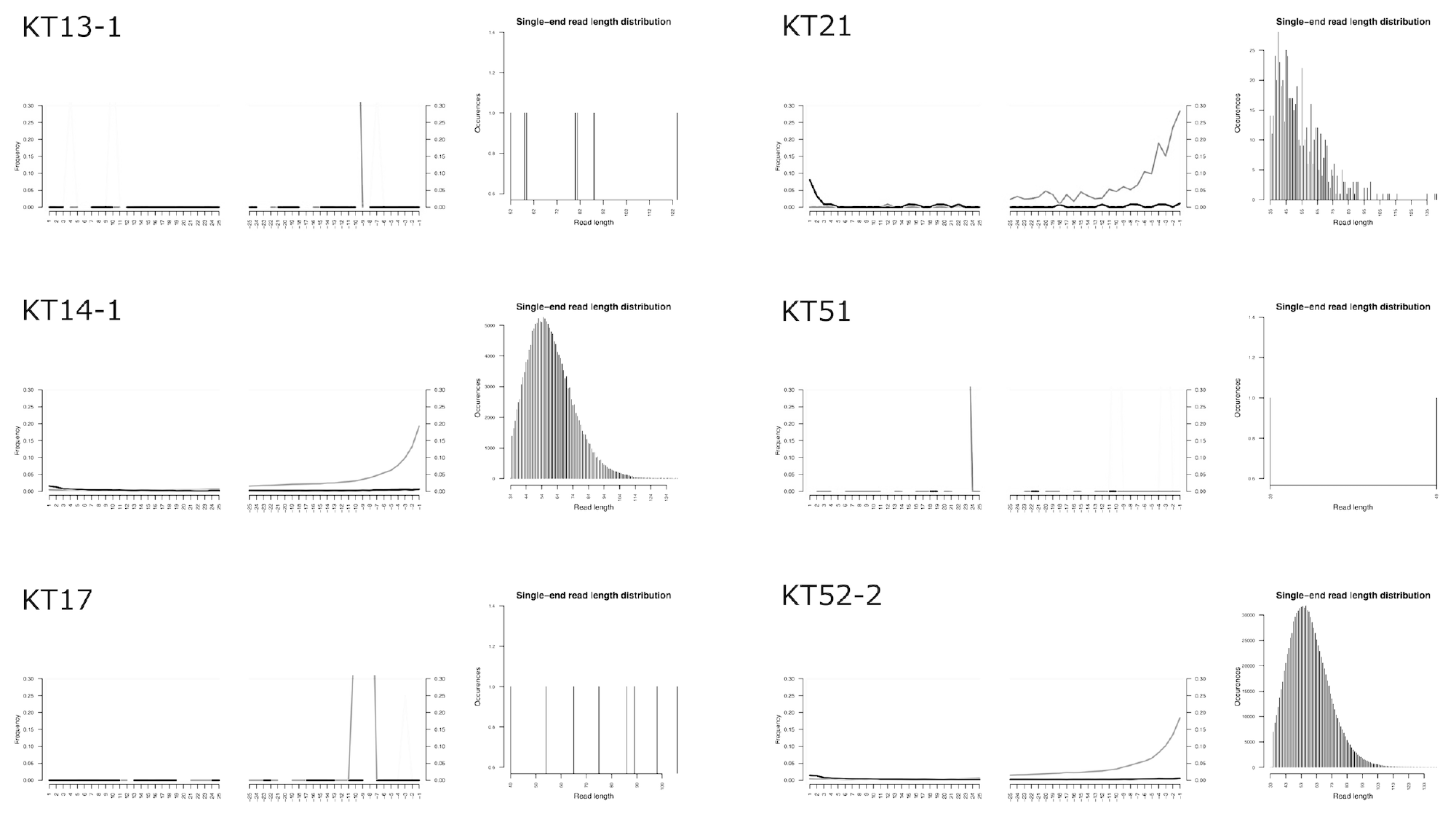

### Slide 7
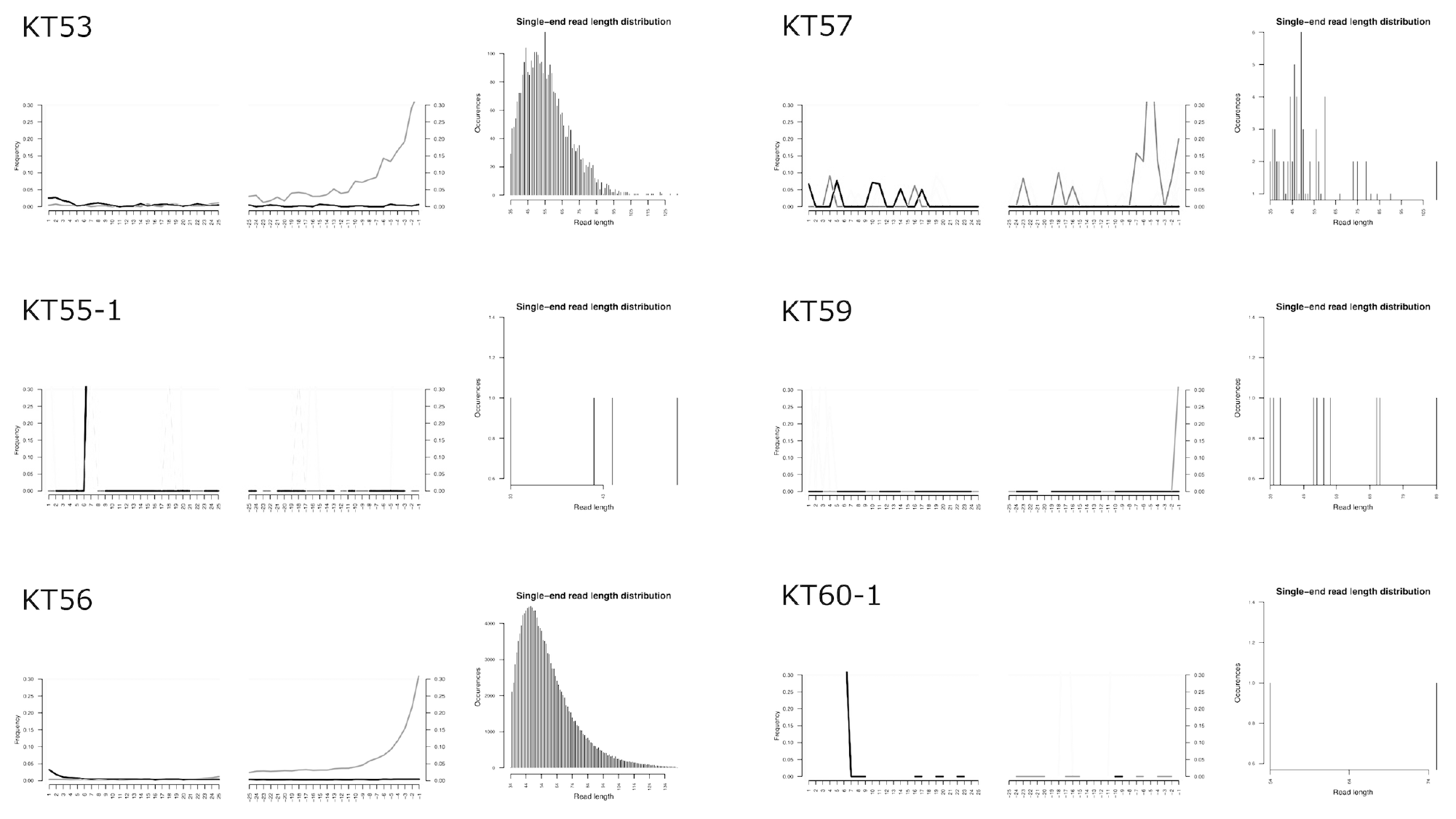

### Slide 8
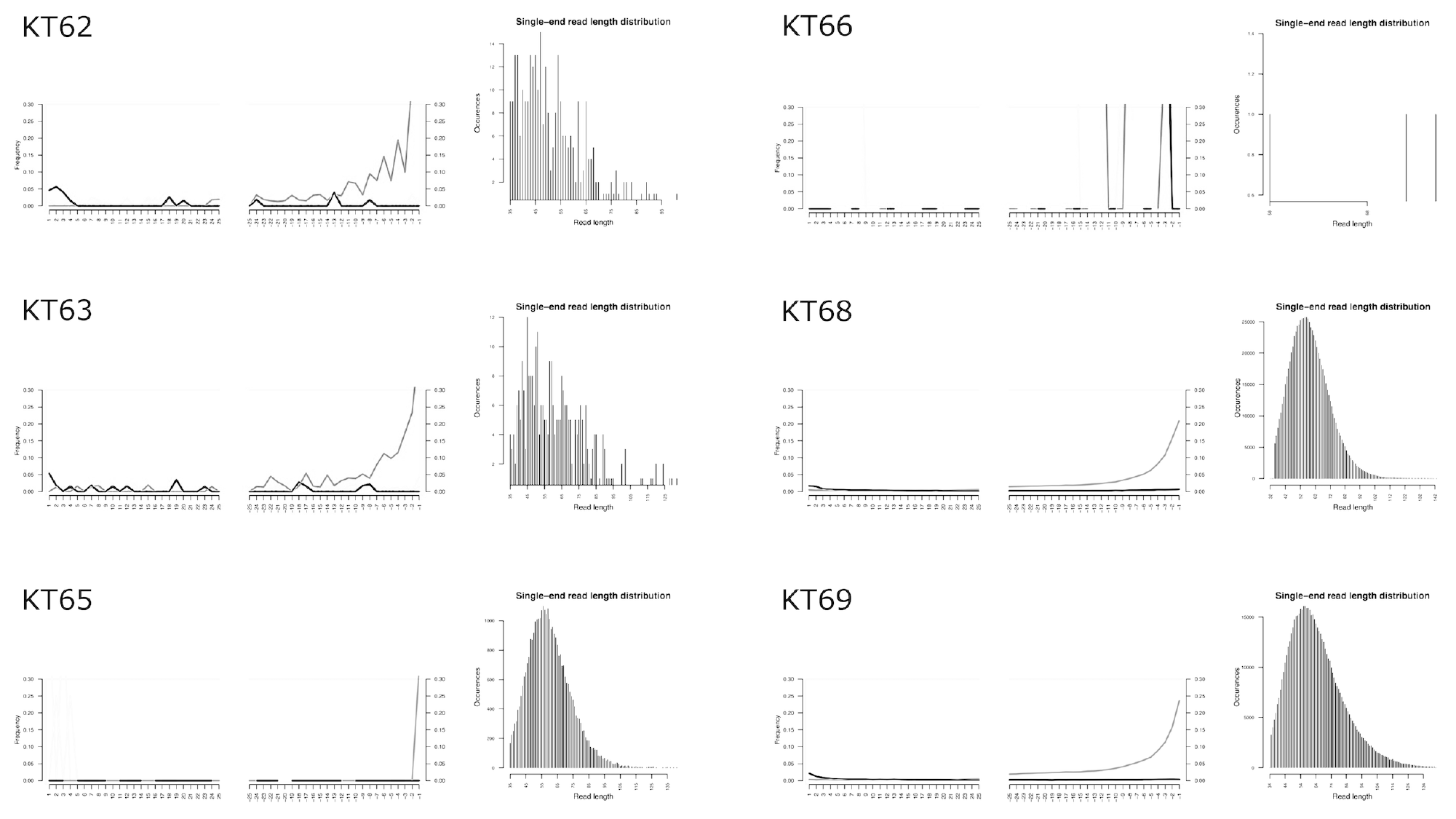

### Slide 9
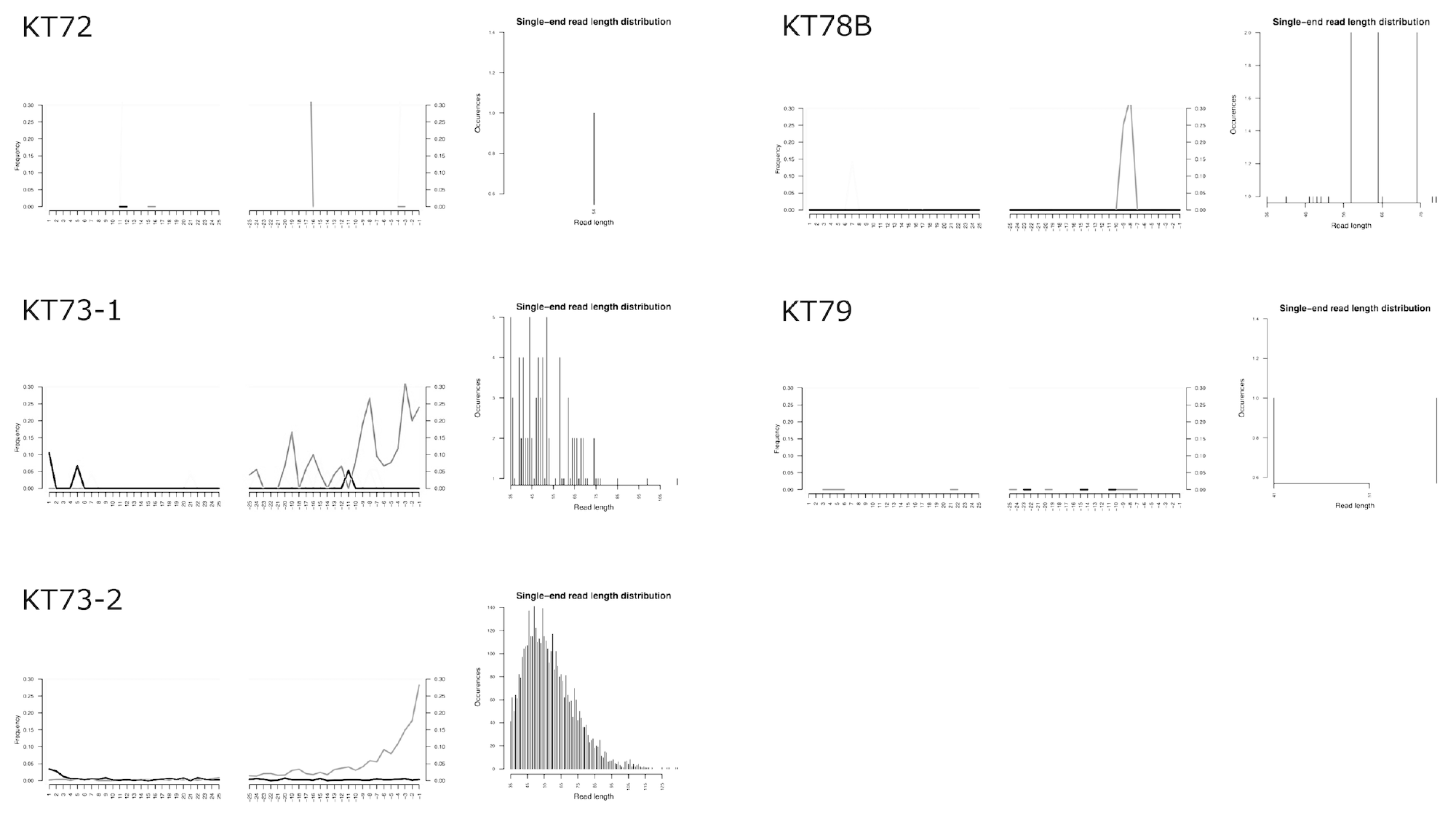

### Slide 10
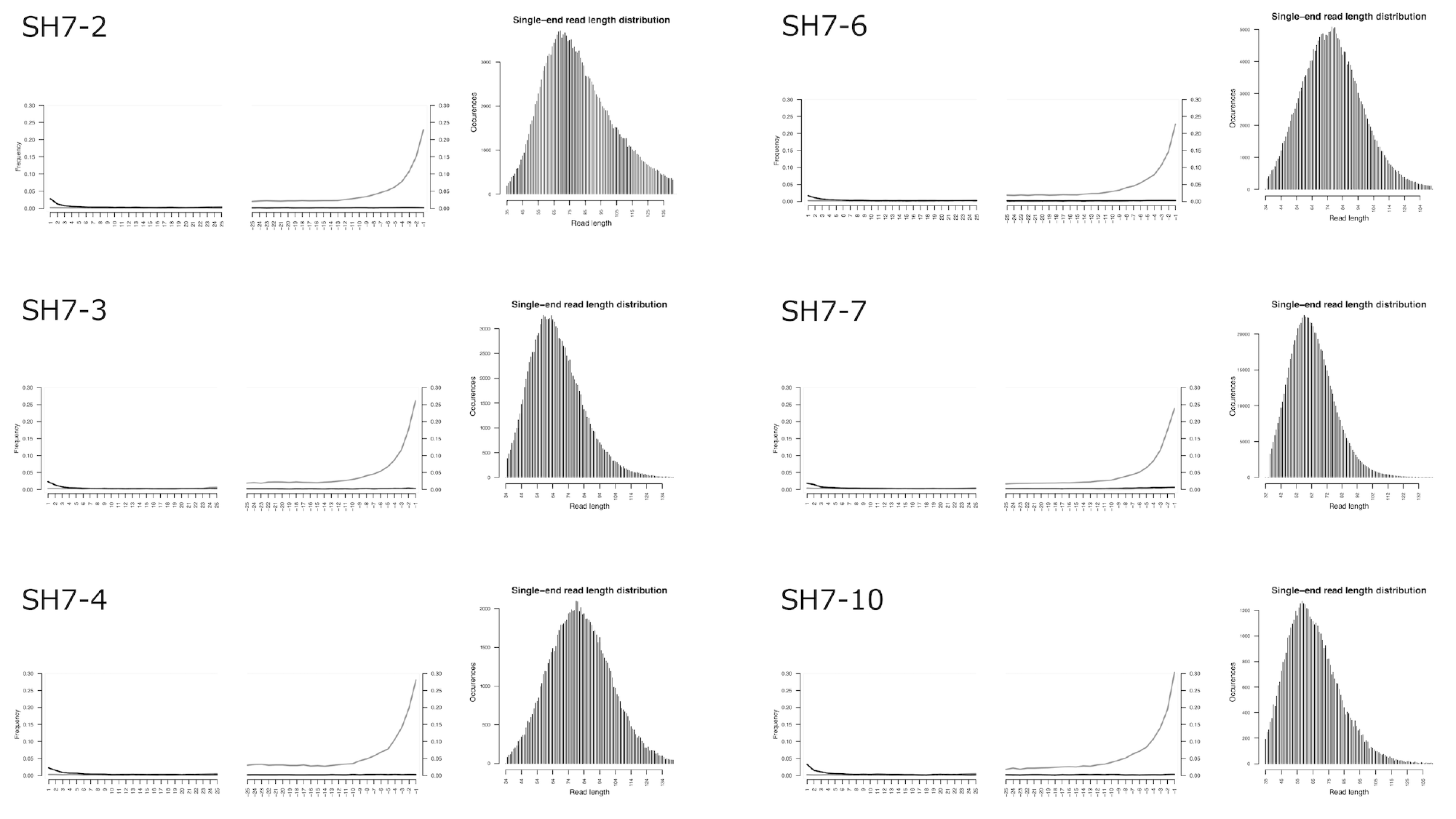

### Slide 11
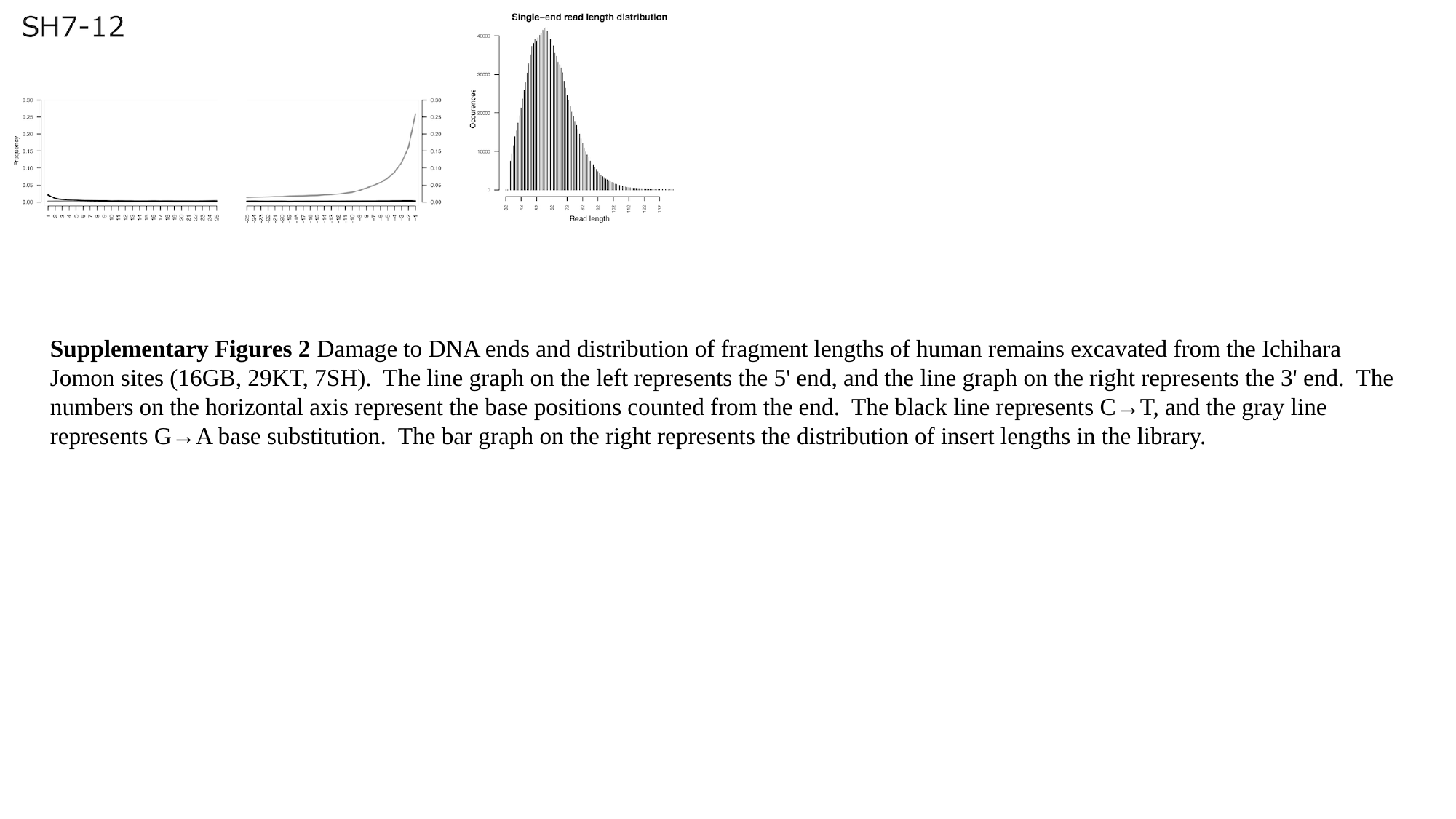

Supplementary Figures 2 Damage to DNA ends and distribution of fragment lengths of human remains excavated from the Ichihara Jomon sites (16GB, 29KT, 7SH). The line graph on the left represents the 5' end, and the line graph on the right represents the 3' end. The numbers on the horizontal axis represent the base positions counted from the end. The black line represents C→T, and the gray line represents G→A base substitution. The bar graph on the right represents the distribution of insert lengths in the library.

### Slide 12
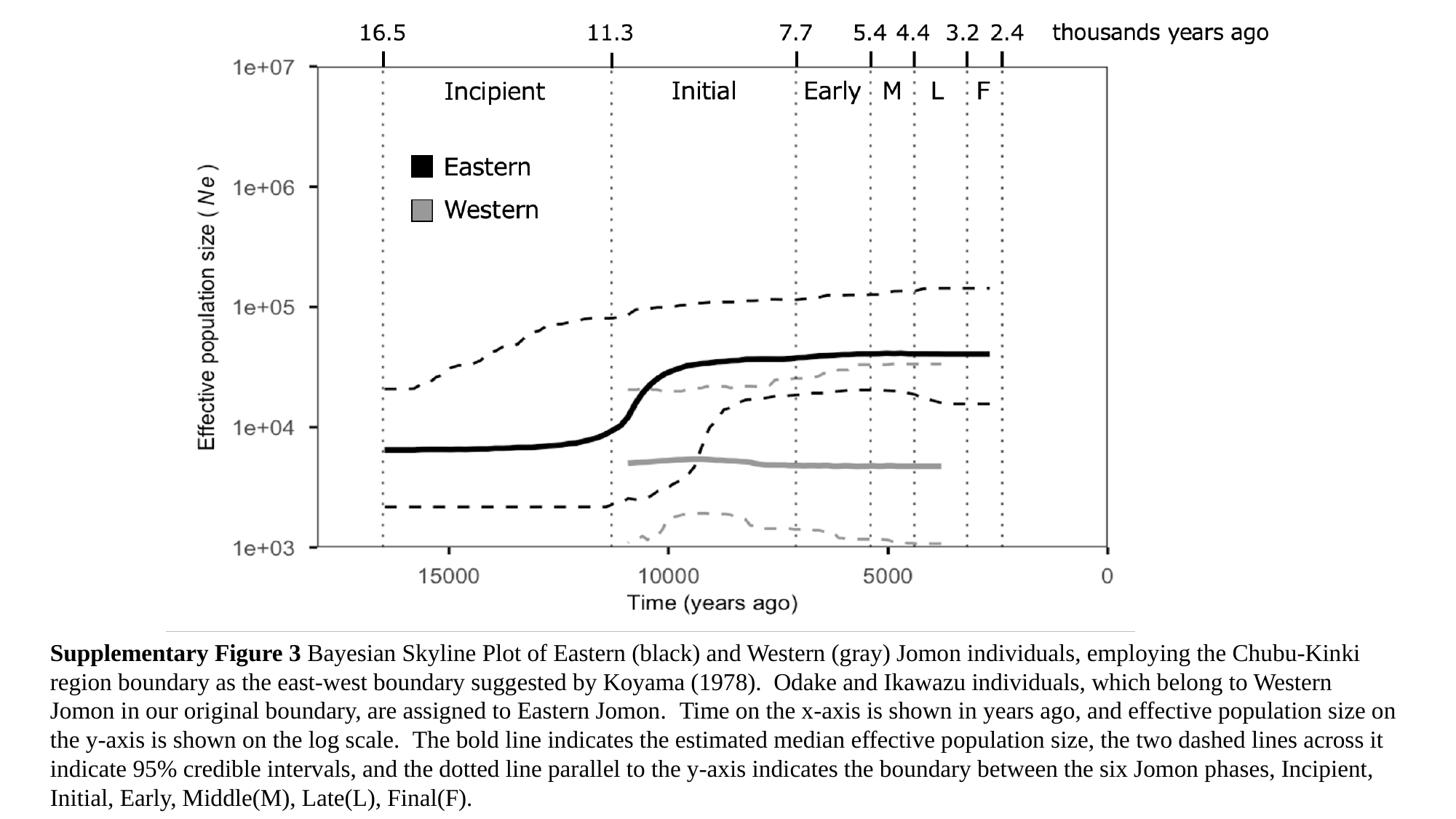

Supplementary Figure 3 Bayesian Skyline Plot of Eastern (black) and Western (gray) Jomon individuals, employing the Chubu-Kinki region boundary as the east-west boundary suggested by Koyama (1978). Odake and Ikawazu individuals, which belong to Western Jomon in our original boundary, are assigned to Eastern Jomon. Time on the x-axis is shown in years ago, and effective population size on the y-axis is shown on the log scale. The bold line indicates the estimated median effective population size, the two dashed lines across it indicate 95% credible intervals, and the dotted line parallel to the y-axis indicates the boundary between the six Jomon phases, Incipient, Initial, Early, Middle(M), Late(L), Final(F).

### Slide 13
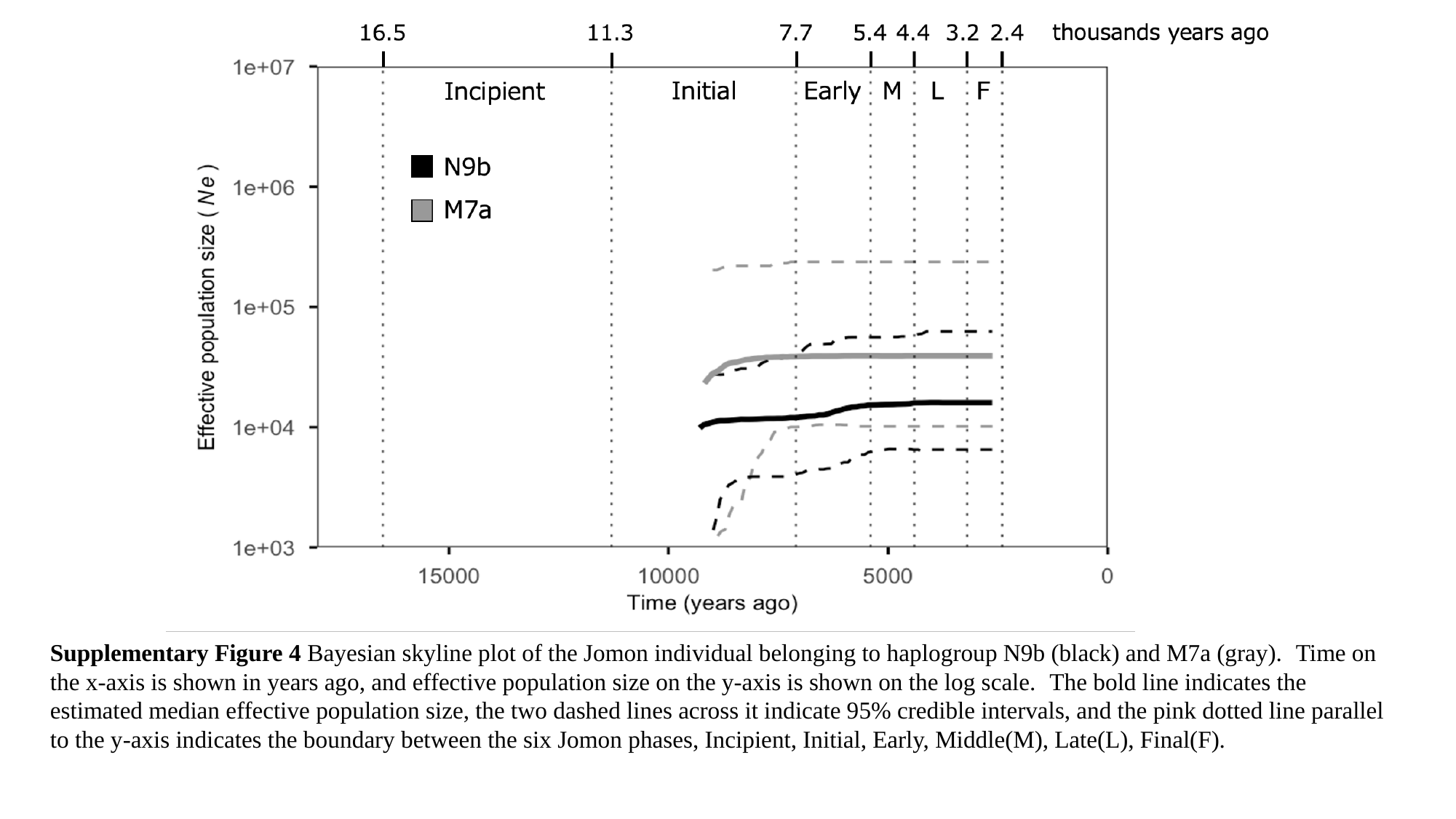

Supplementary Figure 4 Bayesian skyline plot of the Jomon individual belonging to haplogroup N9b (black) and M7a (gray). Time on the x-axis is shown in years ago, and effective population size on the y-axis is shown on the log scale. The bold line indicates the estimated median effective population size, the two dashed lines across it indicate 95% credible intervals, and the pink dotted line parallel to the y-axis indicates the boundary between the six Jomon phases, Incipient, Initial, Early, Middle(M), Late(L), Final(F).
